# A wheat tandem kinase and NLR pair confers resistance to multiple fungal pathogens

**DOI:** 10.1101/2024.08.30.610185

**Authors:** Ping Lu, Gaohua Zhang, Jing Li, Zhen Gong, Gaojie Wang, Lingli Dong, Huaizhi Zhang, Guanghao Guo, Min Su, Yueming Wang, Keyu Zhu, Qiuhong Wu, Yongxing Chen, Miaomiao Li, Baoge Huang, Beibei Li, Wenling Li, Lei Dong, Yikun Hou, Xuejia Cui, Hongkui Fu, Dan Qiu, Chengguo Yuan, Hongjie Li, Jianmin Zhou, Guan-Zhu Han, Yuhang Chen, Zhiyong Liu

**Affiliations:** State Key Laboratory of Plant Cell and Chromosome Engineering, Institute of Genetics and Developmental Biology, The Innovative Academy of Seed Design, Chinese Academy of Sciences, Beijing 100101, China; State Key Laboratory of Molecular Developmental Biology, Institute of Genetics and Developmental Biology, Chinese Academy of Sciences, Beijing 100101, China; College of Bioscience and Resources Environment, Beijing University of Agriculture, Beijing 102206, China; College of Life Sciences, Nanjing Normal University, Nanjing, Jiangsu 210023, China; Institute of Biotechnology, Xianghu Laboratory, Hangzhou, Zhejiang 311231, China; College of Advanced Agricultural Sciences, University of Chinese Academy of Sciences, Beijing 101408, China; Hebei Gaoyi Stock Seeds Farm, Gaoyi, Hebei 051330, China; Yazhouwan National Laboratory, No. 8 Huanjin Road, Yazhou District, Sanya City, Hainan 572024, China; Hainan Seed Industry Laboratory, Sanya City, Hainan 572024, China

## Abstract

Recently discovered tandem kinase proteins (TKPs) are pivotal to the innate immune systems of cereal plants, yet how they initiate plant immune responses remains unclear. This report identifies the wheat protein WTN1, a non-canonical NLR receptor featuring tandem NB-ARC domains, as crucial for *WTK3-*mediated disease resistance. Both WTK3 and its allelic variant Rwt4, known for conferring resistance to wheat powdery mildew and blast respectively, are capable of recognizing the blast effector PWT4, and activate WTN1 to form calcium-permeable channels, akin to ZAR1 and Sr35. This study unveils a unique plant defense mechanism wherein TKPs and associated NLRs operate as “sensor-executor” pairs against fungal pathogens. Additionally, evolutionary analyses reveal a co-evolutionary trajectory of the TKP-NLR module, highlighting their synergistic role in triggering plant immunity.

**One Sentence Summary:** An ancient synergistic TKP-NLR pair triggers innate immunity for multiple disease resistance in wheat.

---

Plant-pathogen interactions can lead to the development of diseases in a susceptible host. To combat pathogens, plants have evolved a complex immune system that encompasses receptors located on the cell surface alongside intracellular receptors. The majority of cloned resistance genes encode for intracellular nucleotide-binding leucine-rich repeat (NLR) receptors (*1*, *2*). Plant NLRs function as singletons, pairs, or within complex networks to bolster plant immunity. Singleton NLRs usually trigger hypersensitive immunity in the host by detecting effectors either directly or indirectly (*3–6*). NLR pairs consist of a sensor NLR recognizing effectors and a helper NLR engaged in complex formation with the sensor NLR (*3, 7*). These paired NLRs arrange in a head-to-head orientation in the genomes and undergo hetero-oligomerization, forming distinctive “dimers of heterodimers” structures that lead to the initiation of an immune response and cell death (*7*). Within the NLR network, helper NLRs are members of the NLR required for cell death (NRC) family, displaying distinct specificities towards varying sensor NLRs (*3, 8*).

NLR proteins can aggregate into a sophisticated oligomeric structure known as the resistosome when they recognize pathogen effectors (*9,10*). The wheat stem rust resistance gene *Sr35,* derived from einkorn wheat (*Triticum monococcum*), encodes an NLR receptor that directly engages with the avirulence (Avr) effector AvrSr35. This interaction results in the formation of a pentameric Sr35-AvrSr35 complex, thereby imparting resistance against *Puccinia graminis* f. sp. *tritici* (*11*). Additionally, the *Arabidopsis* NLR protein ZAR1 is found to pre-associate in a complex with the pseudokinase RKS1 to detect the uridylyltransferase effector AvrAC of *Xanthomonas campestris*. The PBL2 kinase is modified by AvrAC and subsequently recruited to the ZAR1-RKS1 complex, culminating in the formation of a resistosome that functions as a calcium channel (*6,9*). The recent elucidation of both inactive and activated structures of ZAR1 and Sr35 has significantly advanced our understanding of the operational mechanisms of plant NLRs, shedding light on the intricate molecular interactions underpinning plant immunity (*6, 9–11*).

Recent discoveries within the *Triticeae* tribe have highlighted the significance of tandem kinase proteins (TKPs) as an essential family of intracellular proteins that can contribute to plant immune responses (*12*). This family encompasses a series of wheat disease resistance genes against stripe rust (*Yr15*/*WTK1*) (*13*), stem rust (*Sr60*/*WTK2* and *Sr62*/*WTK5*) (*14*, *15*), leaf rust (*Lr9*/*WTK6-vWA*) (*16*), powdery mildew (*Pm24*/*WTK3* and *WTK4*) (*17*, *18*), wheat blast *Rwt4* (*19*), along with the barley stem rust resistance gene *Rpg1* (*20*). Despite the identification of these genes, the precise mechanisms by which TKPs mediate immune responses remain elusive.

In this study, we report a wheat protein WTN1, an atypical NLR with tandem NB-ARC domains, cooperates with WTK3/Rwt4 to constitute a TKP-NLR immune signaling pathway. Upon sensing the pathogen Avr PWT4, this TKP-NLR pair is activated to form a calcium-permeable channel, thereby enhancing resistance to fungal pathogens in wheat.

## A non-canonical NLR is required for *WTK3*-mediated powdery mildew resistance

To identify genes implicated in the *WTK3-*mediated powdery mildew resistance pathway, we analyzed 3,860 M_2_ progenies of EMS-mutagenized wheat landrace Hulutou (HLT), inoculated with the *Blumeria graminis* f. sp. *tritici* (*Bgt*) isolate E09, an avirulent isolate to *Pm24* (*WTK3*). Among the susceptible mutants obtained, 16 without mutations in *WTK3* were selected for subsequent RNA-sequencing and PCR amplification. Mutations leading to amino acid substitutions were identified in five mutants (M1048^L122F^, M116^G408E^, M1037^T528I^, M1136^G589S^, and M189^D636N^) at an atypical NLR protein featuring two NB-ARC domains (NBD) (fig. S1, S2A, C), hereafter referred to as Wheat Tandem NBD 1 (WTN1), located 114 kb adjacent to *WTK3* on chromosome 1DS (Fig. 1, A and B). To test whether *WTN1* is required for the *WTK3*-mediated powdery mildew resistance, crosses were made between two *WTK3* nonsense mutants, M410^W429*^ and M1091^W478*^ (*17*), and the five *WTN1* mutants (Fig. 1, A and B, fig. S2B). All F_1_ plants from the ten crosses exhibited resistance to *Bgt* isolate E09 (Fig. 1B). Conversely, F_1_ plants from half diallel crosses among the *WTN1* mutants showed high susceptibility (fig. S3), demonstrating the essential role of *WTN1* in *WTK3-*dependent resistance to *Bgt* pathogen.

**Fig. 1.**
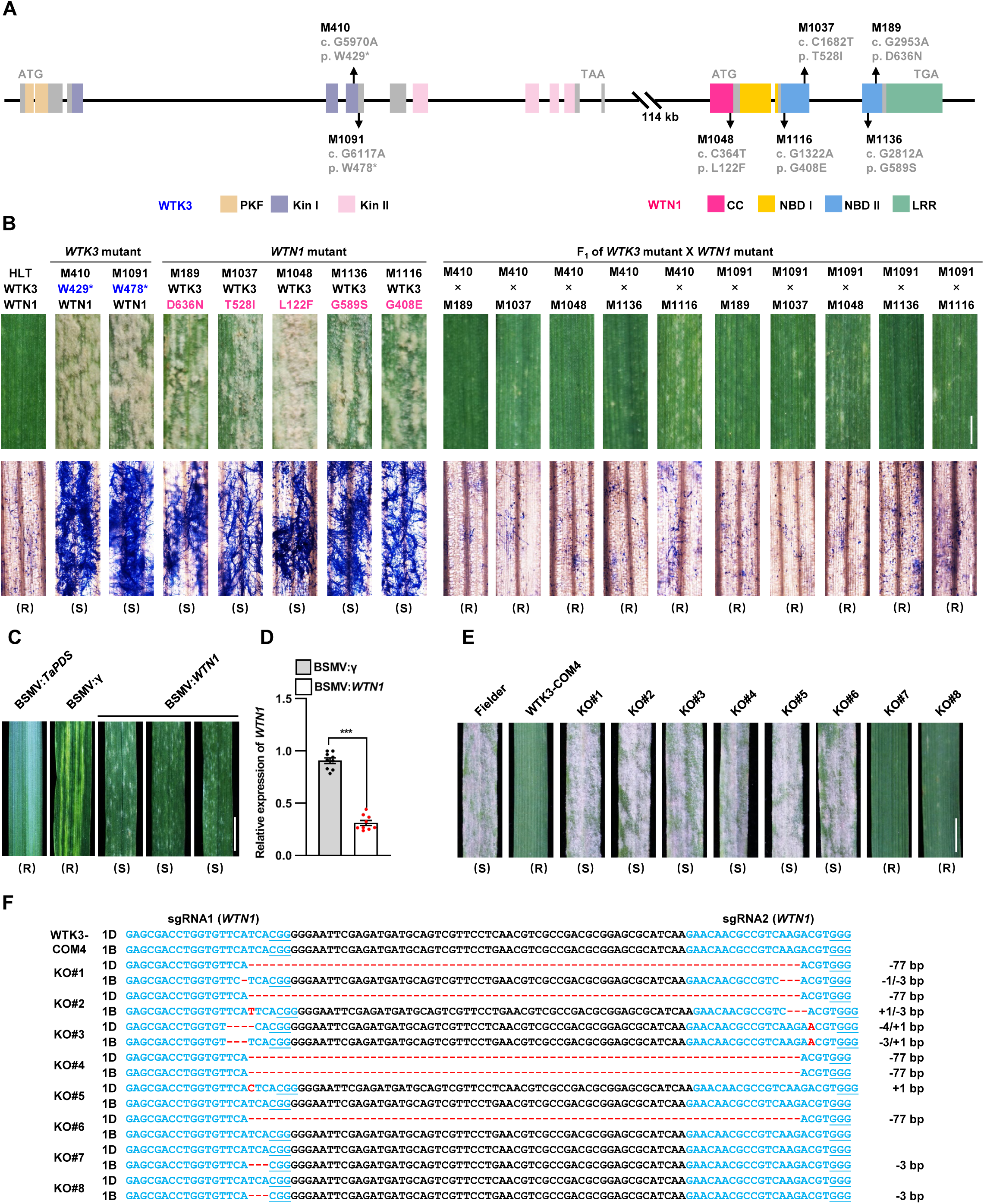
Functional roles of TKP *WTK3* and NLR *WTN1* in powdery mildew resistance at the *Pm24* locus. **(A)** Chromosome location and gene structures of TKP *WTK3* and NLR *WTN1* on chromosome 1DS, including mutations (marked in black) and their impacts on the translated proteins in susceptible mutant lines. Gene components are depicted as follows: exons (Boxes) and introns (lines). The functional domains of WTK3, including pseudo-kinase fragment (PKF), Kinase I (Kin I), Kinase II (Kin II), and those of WTN1, including coiled-coil (CC), NB-ARC I (NBD I), NB-ARC II (NBD II), and leucine-rich repeat (LRR), are color-coded in light-orange, light-blue, light-pink, red, orange, blue, and green, respectively. **(B)** Comparison of powdery mildew infection on leaves of the wild-type HLT, susceptible mutants of *WTK3* and *WTN1*, and their F_1_ hybrids with *Blumeria graminis* f. sp. *tritici* (*Bgt*) isolate E09 at 14 days post-inoculation (dpi). Images show representative leaves (Scale bar, 0.3 cm) and Trypan blue staining to highlight fungal structures (Scale bar, 200 μm). The wild type genotypes of *WTK3* and *WTN1* in the HLT mutants are marked in black, the mutant genotypes of *WTK3* and *WTN1* are marked in blue and red, respectively. R indicated the plants resistance to powdery mildew, S indicated the plants susceptibility to powdery mildew. **(C)** Disease symptoms on the third leaves of HLT plants inoculated with *Bgt* E09 at 14 dpi, following pre-inoculation with BSMV:*TaPDS* or BSMV:γ or BSMV:*WTN1* (Scale bar, 0.5 cm). **(D)** Expression levels of *WTN1* in BSMV:γ and BSMV:*WTN1* plants inoculated with *Bgt* isolate E09 at 14 dpi. Data are expressed as means ± SE from three independent experiments, analyzed using Student’s *t*-test (*P* < 0.001). **(E)** Infection phenotypes of *WTN1* CRISPR/Cas9 knockout (KO) mutants. Detached leaves of Fielder, *WTK3* transgenic line WTK3-COM4, and eight different CRISPR/Cas9 KO events (KO#1 - KO#8) were inoculated with *Bgt* isolate E09 (Scale bar, 0.5 cm). **(F)** Genotypes of *WTN1* and *WTN-1B* in WTK3-COM4 and the eight CRISPR/Cas9 KO mutants. sgRNA1 and sgRNA2 targeted both the *WTN1* and *WTN-1B* homoeolog, with the sgRNA sites marked in blue and mutations highlighted in red.

To further determine the role of *WTN1* in *WTK3*-mediated powdery mildew resistance, *WTN1* expression was suppressed in the resistant HLT using *barley stripe mosaic virus* (BSMV)-induced gene silencing (VIGS). The expression of *WTN1* was significantly reduced, leading to increased production of powdery mildew spores in the *WTN1* knockdown plants compared to the BSMV-γ control plants at 14 days post-inoculation (dpi) with *Bgt* isolate E09 (Fig. 1, C and D). The findings demonstrate that silencing *WTN1* compromises *WTK3-*mediated defense against *Bgt*.

We then developed *WTN1* CRISPR/Cas9 knockout mutants in a previously generated transgenic Fielder line, WTK3-COM4, which carries a *WTK3* transgene and is resistant to *Bgt* (*17*). Wild-type Fielder contains a *wtk3* susceptible allele, a functional *WTN1* allele on 1DS, and a homoeolog, *WTN1-1B,* on 1BS that has 91.66% amino acid sequence identity to *WTN1* (fig. S4). Two specific guide RNAs (gRNA1 and gRNA2) targeting the 5-terminus of exon 1 of *WTN1* and *WTN1*-*1B* were introduced into WTK3-COM4 using Agrobacterium-mediated transformation (fig. S4). This approach yielded eight independent knockout mutants, with KO#1 to KO#4 being double mutants for *WTN1* and *WTN1*-*1B*, KO#5 and KO#6 only affecting *WTN1*, KO#7 and KO#8 specifically targeting *WTN1*-*1B* (Fig. 1, E and F). When inoculated with *Bgt* isolate E09, KO#1 to KO#6 displayed high susceptibility to powdery mildew, in contrast to the high resistance observed in KO#7 and KO#8, as well as in the non-edited control (Fig. 1E). These findings indicate that *WTN1* is indispensable for *WTK3*-mediated powdery mildew resistance.

## WTK3 interacts with WTN1

Upon discovering that WTN1 is integral to WTK3 function, we conducted a comprehensive analysis to characterize their interaction. Bimolecular fluorescence complementation (BiFC) assay yielded YFP fluorescence upon co-expression of WTK3:YFP^C^ with WTN1:YFP^N^, indicating a direct *in vivo* WTN1-WTK3 interaction (Fig. 2A). This interaction is further confirmed by split firefly luciferase complementation (SFLC) assay in *Nicotiana benthamiana* leaves (Fig. 2B) and co-immunoprecipitation (Co-IP) assay in protoplasts isolated from the wheat cultivar Fielder (Fig. 2C). Moreover, additional Co-IP assay implicated the C-terminal Kin II domain of WTK3 (residues 521 to 893) as the specific region mediating interaction with WTN1 (Fig. 2D). These results demonstrate a direct interaction between WTK3 and WTN1, suggesting their synergistic role in plant immune regulation.

**Fig. 2.**
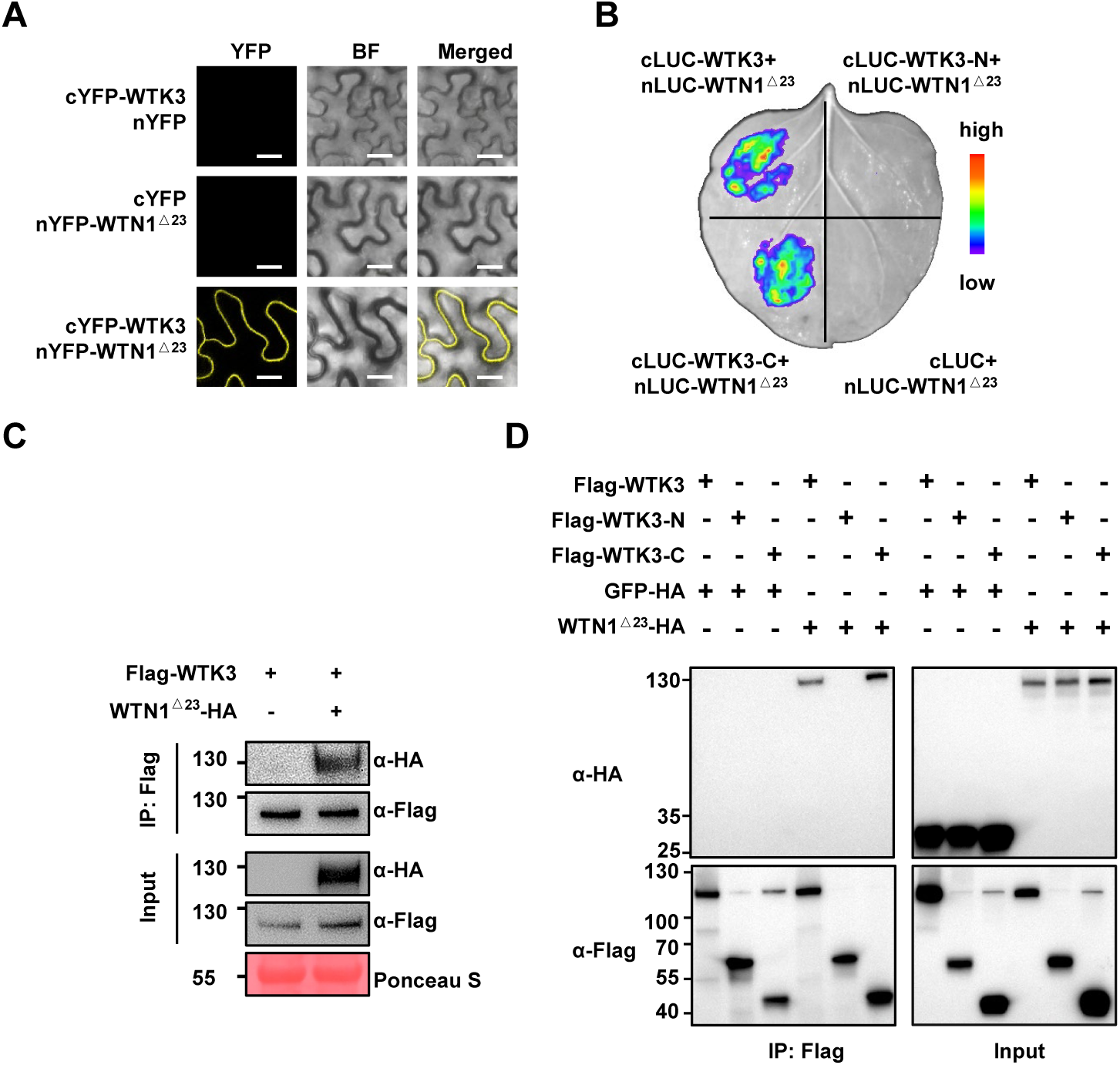
WTN1 interacts with WTK3. **(A)** Bimolecular fluorescence complementation (BiFC) assay showing *in vivo* interaction between WTK3 and WTN1^Δ 23^ (WTN1^24-1038^) in *Nicotiana benthamiana* leaves. WTK3 was fused to the C-terminal half of yellow fluorescence (cYFP), and WTN1 ^Δ 23^ to the N-terminal half (nYFP). Fluorescence indicates protein-protein interaction. Scale bar, 20 μm. **(B)** Split firefly luciferase complementation (SFLC) assay demonstrating the interaction between WTK3, WTK3-N^1-500^ and WTK3-C^521-893^ with WTN1 ^Δ 23^(WTN1^24-1038^) in *N*. *benthamiana* leaves. WTK3, WTK3-N^1-500^ and WTK3-C^521-893^ were fused to the C-terminal half of luciferase (cLUC) and WTN1^Δ23^ (WTN1^24-1038^) to the N-terminal half (nLUC), with luminescence indicating interaction. **(C)** Co-immunoprecipitation (Co-IP) assay confirming the interaction between WTK3 and WTN1^Δ23^ in wheat protoplasts from the Fielder variety. WTK3 was tagged with Flag, and WTN1^Δ 23^ with HA epitope. **(D)** Co-IP assay confirming the interaction between WTK3, truncated WTK3 (WTK3-N^1-500^ and WTK3-C^521-893^) and WTN1^Δ23^ (WTN1^24-1038^) in *N*. *benthamiana*, tagged with Flag and HA, respectively.

## RWT4/WTK3 detect avirulent protein and induce WTN1 oligomerization

Next, we sought to unravel the functional role and activation mechanism of WTN1. While identifying its specific Avr protein is crucial, this challenge has been overcome by discovering Rwt4 (an allelic variant of WTK3) along with its effector PWT4 (*19*, *21*). Compared to WTK3, Rwt4 has a two-residue (K400/G401) insertion in a loop within the Kin I domain, caused by a 6-bp InDel (AAAGGA/-) in the fifth exon of the two alleles (Fig. 3A, B, fig. S5A, B). Thus, we took advantage of the Rwt4 together with its cognate effector PWT4 to elucidate how TKP detects pathogen effectors and activates NLR to trigger immune responses.

**Fig. 3.**
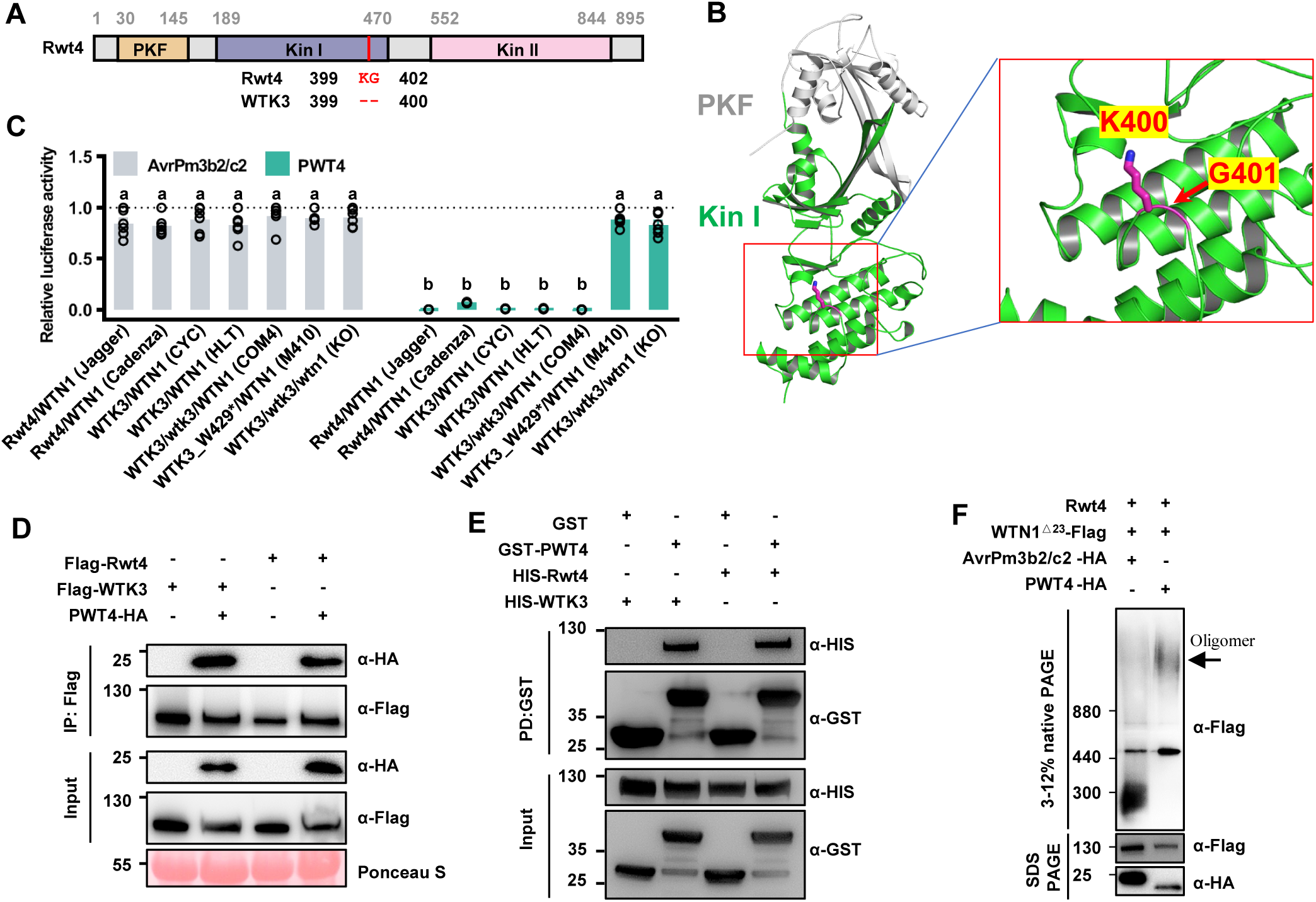
Rwt4 and WTK3 interact with PWT4. **(A)** Comparison of amino acid sequences of Rwt4 and WTK3. Differences in amino acid residues between Rwt4 and WTK3 are marked in red. The pseudo-kinase fragment (PKF), Kinase I (Kin I), and Kinase II (Kin II) domains of Rwt4/WTK3 are color-coded in light-orange, light-blue and light-pink, respectively. **(B)** The N-terminus of Rwt4 protein structures was predicted using AlphaFold2. The positions of K400G401 in Kin I were shown. **(C)** Protoplast assays in various wheat genotypes transfected with PWT4 and AvrPm3b2/c2, comparing genotypes with different combinations of *Rwt4*/*WTN1* and *WTK3*/*WTN1* alleles. Cell death was quantified using relative luminescence. AvrPm3b2/c2 treatment defines the relative baseline (mean ± s.e.m.; n = 6) in protoplasts of different wheat genotypes. Different letters denote the significance tested by one-way analysis of variance (ANOVA) and Tukeys post -hoc test at *P* < 0.05. Exact *P* values are provided in Supplementary Table 5. **(D)** Co-IP assay confirming the interaction between Rwt4/WTK3 and PWT4 in *N*. *benthamiana*, tagged with Flag and HA, respectively. **(E)** GST pull-down assay demonstrating direct protein-protein interactions between Rwt4/WTK3 and PWT4. **(F)** Oligomerization analysis: PWT4 trigger WTN1 oligomerization in wheat Jagger (Rwt4/WTN1) protoplasts, confirmed by blue native PAGE (BN-PAGE), and SDS-PAGE for protein expression, with consistent results across two experiments.

We then evaluated whether PWT4 could be recognized by Rwt4 or WTK3 and activate NLR to induce hypersensitive responses. To mimic pathogen attack, we expressed PWT4 in various wheat protoplasts harboring endogenous Rwt4 or WTK3 and WTN1, and performed luminescence-based cell viability assays. Relative luminescence activity measurements revealed that expression of PWT4 induced severe cell death in protoplasts with Rwt4/WTN1 from Jagger and Cadenza, as well as in those with WTK3/WTN1 from WTK3-COM4, CYC, and HLT lines. Notably, protoplasts possessing only WTN1 or WTK3, such as from WTK3^W429^* or WTN1-KO, did not undergo cell death (Fig. 3C). Control treatment with the irrelevant Avr, AvrPm3bc/c2 (*22*), did not result in cell death, confirming the specificity of PWT4-induced hypersensitive response. These results demonstrate that both TKP and NLR are required to trigger hypersensitive responses, consistent with their synergistic role in disease resistance.

To delineate the underlying mechanism, we further analyzed whether PWT4 could directly interact with RWT4 or WTK3. When Rwt4 or WTK3 was co-expressed with PWT4 in *N. benthamiana* leaves, our Co-IP assay revealed a direct protein-protein interaction between Rwt4 or WTK3 and PWT4 (Fig. 3D), which was further confirmed by an *in vitro* GST pull-down assay (Fig. 3E). These results consistently demonstrate that PWT4 can recognize and directly bind to both Rwt4 and WTK3. This is not surprising, considering that WTK3 and Rwt4 only differ by two residues in Kin I. However, this finding also raises the possibility that the *WTK3* allele may also confer resistance to wheat blast.

In cell death assays conducted in *N. benthamiana*, individual expression of Rwt4, WTK3, WTN1, or PWT4 did not induce any discernible hypersensitive responses (HR) (fig. S6A). Co-expression of WTK3 or Rwt4 with PWT4, or WTN1 with PWT4, also failed to trigger HR. However, simultaneous expression of both Rwt4/WTK3 and WTN1, with or without PWT4, resulted in HR manifestation within 3 days (fig. S6B).

Truncations in the N-terminal α1 helix of WTN1 (including WTN1^Δ6^, WTN1^Δ10^ and WTN1^Δ23^), along with deletions of NB-ARC, led to abolition of cell death activity, confirming its essential role in triggering cell death (fig. S6C). Furthermore, co-expression of WTN1^Δ23^ alongside Rwt4 and PWT4 in wheat protoplasts resulted in noticeable oligomerization of WTN1^Δ23^, as revealed by BN-PAGE separation of total proteins (Fig. 3F), contrasting with the control group expressing irrelevant AvrPm3bc/c2.

## WTN1 activation forms calcium-permeable channels for triggering immunity

To test the functional role of the activated WTN1, we co-expressed WTN1 with PWT4, Rwt4 or WTK3 in *Xenopus laevis* oocytes, and performed two-electrode voltage clamp (TEVC) measurements (Fig. 4A and B). No detectable currents were produced when WTN1, PWT4, Rwt4, and WTK3 cRNAs were injected individually or in pairs. However, a dramatic shift was observed when WTN1 was co-expressed with both PWT4 and Rwt4 (or WTK3), leading to the emergence of substantial currents, suggesting the formation of resistosome channels reminiscent of ZAR1 and Sr35 (*9*, *11*). Given that ZAR1- and Sr35-dependent currents recorded from *Xenopus* oocytes were confounded by Cl^-^ currents (elicited by Ca^2+^ influx) mediated through the endogenous calcium-activated chloride channel (CaCC) (*6*, *9*, *11*), it is plausible to hypothesize that WTN1-dependent currents might have a similar component. Indeed, these currents were partially diminished upon applying a CaCC inhibitor (CaCCinh-A01) and completely counteracted by the calcium channel blocker LaCl_3_ (Fig. 4C), indicating these currents as a proxy for Ca^2+^ entry via the WTN1 channel.

**Fig. 4.**
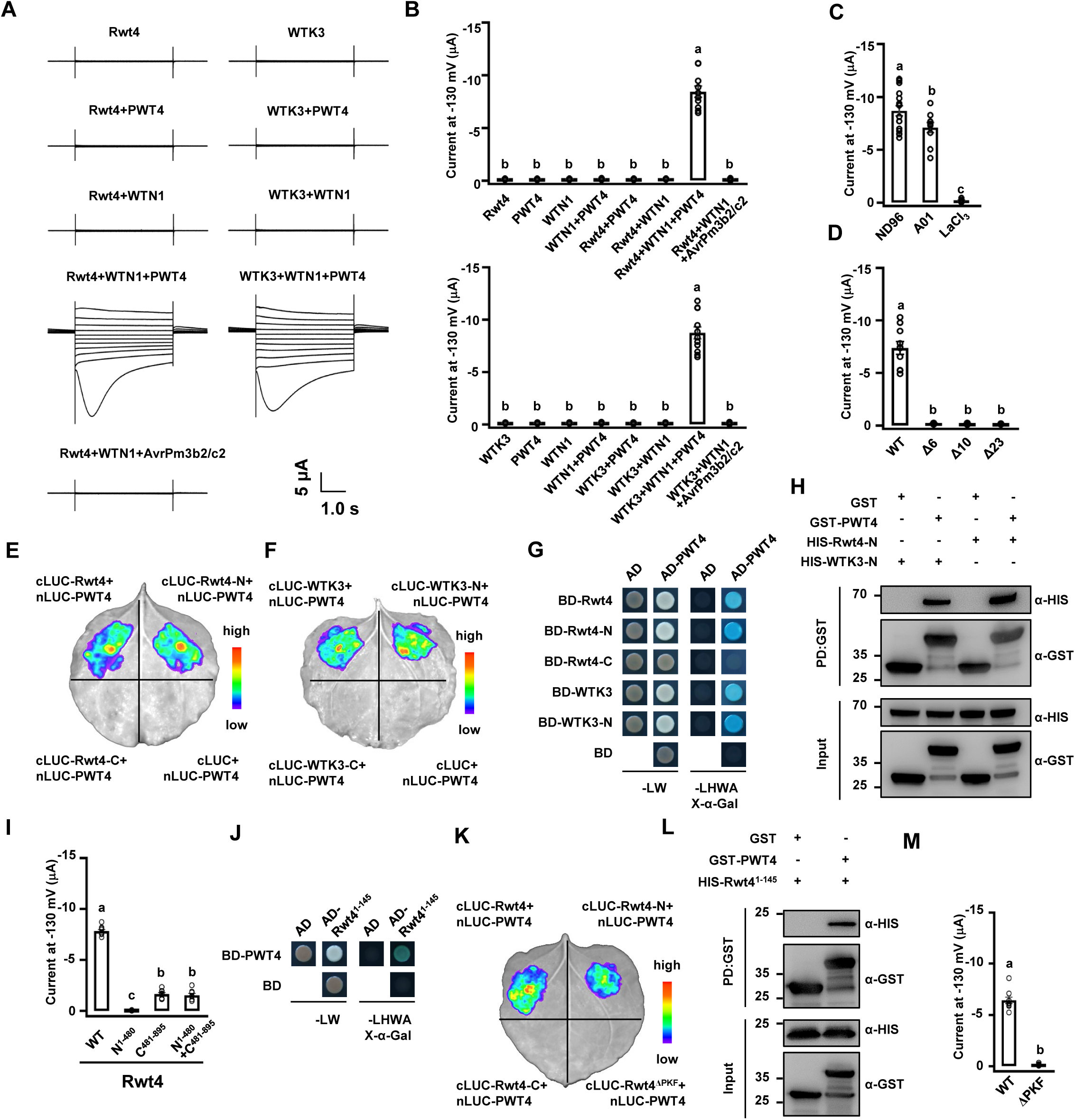
Rwt4/WTK3 sense PWT4 to activate WTN1 for forming calcium-permeable channels. **(A and B)** Two-electrode voltage clamp (TEVC) measurements in *Xenopus* oocytes expressing Rwt4/WTN1/PWT4 and WTK3/WTN1/PWT4, with control oocytes injected with Rwt4/WTN1/AvrPm3b2/c2. TEVC recordings were performed in ND96 solution. Representative current traces over arrange of voltages (from -130 mV to +70 mV in 20 mV increment) are shown in **(A)**, and current amplitudes measured at -130 mV are shown in **(B)**. **(C)** Electrophysiological effects of inhibitors: The impact of CaCCinh-A01 (Ca^2+^-activated chloride channel inhibitor) and LaCl_3_ (Ca^2+^ channel blocker) on the WTN1-dependant currents in ND96 solution, showing pre-incubation effects on WTK3/WTN1/PWT4-injected oocytes. **(D)** Truncational analysis of the α1-helix at the N-terminus of WTN1. **(E - H)** Interaction of N-terminus of Rwt4/WTK3 with PWT4 detected by split luciferase complementation (SFLC) **(E and F)**, Yeast two-hybrid (Y2H) **(G)** and GST pull-down **(H)** assays. **(I)** Truncational analysis of the N-terminus and C-terminus of Rwt4. **(J-M)** Interaction of PKF domain of Rwt4 with PWT4 detected by Y2H **(J)**, SFLC **(M),** GST pull-down **(L)** assays. **(M)** Truncational analysis of the PKF of Rwt4. Data are presented as means ± SEM, n ≥ 8. Different letters denote the significance tested by one-way analysis of variance (ANOVA) and Tukeys post -hoc test at *P* < 0.01. Exact *P* values are provided in Supplementary Table 6.

The role of activated WTN1 in ion channel formation was also investigated through N-terminal truncations and EMS-induced mutations. N-terminal deletions (Δ6, Δ10, and Δ23), which did not induce cell death in *N. benthamiana* (fig. S6C), suppressed the WTN1-dependent currents (Fig. 4D), thereby highlighting the pivotal role of the N-terminal α1 helix in forming channel assembly capable of penetrating the cell membrane. Among the five EMS-induced WTN1 mutations, four (L122F, G408E, T528I, and G589S) led to a substantial decrease in current, rendering it virtually undetectable, while D636N only partially impaired it (fig. S7A). Predictive modeling through Alphafold2 has elucidated that the amino acid substitutions in mutants WTN1^G408E^, WTN1^T528I^, WTN1^G589S^, and WTN1^D636N^ are located either within or proximate to the ATP-binding site (fig. S7B). This site is integral to the assembly of the resistosome, a complex crucial for NLR-mediated immune responses (*10*). These findings reinforce the notion that activated WTN1 mediates plant immunity via channel activity, whereas loss-of-function (LOF) mutations disrupt its function by interfering with this mechanism.

While the C-terminal segment of TKP binds NLR, its N-terminal portion is primarily responsible for Avr detection. Our results from *in vivo* SFLC (Fig. 4E and F) and Y2H (Fig. 4G), as well as *in vitro* GST-pull down (Fig. 4H), demonstrate that the N-terminal portions of both Rwt4 (Rwt4-N) and WTK3 (WTK3-N) directly interact with PWT4. Further electrophysiological studies revealed that the N-terminal segment alone failed to stimulate currents, while the C-terminal portion alone or their combination elicited only small currents (Fig. 4I). Notably, five previously identified LOF mutations in WTK3, two at the N-terminus (E372K and G389S) and three at the C-terminus (A600T, G718R, and P835S), reduced channel activity, ranging from moderate to negligible levels (fig. S7C). These findings suggest that both kinase domains and their structural integrity are essential for Avr detection, NLR activation, and resistosome channel assembly.

It was further discovered through *in vivo* SFLC (Fig. 4K) and *in vitro* GST-pull down (Fig. 4L) that the PKF (residue 1-145) region in Rwt4 is capable of directly binding to PWT4, demonstrating its critical role in Avr detection. Electrophysiological study has further established that Rwt4 mutant lacking the PKF exhibits virtually no current, indicating an inability to recognize Avr (Fig. 4M). This deficiency likely prevents the activation of NLR, which is essential for forming the ion channels that confer disease resistance.

## Rwt4/WTK3 and WTN1 underwent co-evolution

Rwt4/WTK3 contains a degenerated PKF and two intact kinase domains, Kin I and Kin II (fig. S8A). Kin I and Kin II were found broadly across the Pooideae subfamily, whereas the PKF is specific to the Triticeae tribe and *Lolium perenne* (fig. S9). Phylogenetic analysis delineated PKF/Kin I and Kin II into two distinct clades, with PKF sequences clustering within the diversity of Kin I sequences (fig. S8B-D). These results suggest that Rwt4/WTK3 may have arisen from the fusion of two distinct kinase domains prior to the last common ancestor (LCA) of Pooideae, with the PKF emerging from a recent duplication of Kin I in the early evolution of Triticeae. Critical kinase catalytic motifs, including the ATP-binding, HRD, and DFG motifs, are well conserved in Kin I but absent or not conserved in Kin II and the degenerated PKF (fig. S8A), indicating pseudo-kinase nature of PKF and Kin II. The ratio of nonsynonymous to synonymous nucleotide substitution rates (dN/dS) for PKF (0.964) and Kin I (0.896) was markedly higher than that for Kin II (0.487), suggesting that PKF and Kin I have undergone a greater degree of positive selection, potentially driven by host-pathogen interactions (fig. S8E). Moreover, several residues crucial to immune function (such as G389, W429, A600, G679, G718, R834, and P835) (*17*) were found to be under negative selection, further emphasizing their importance in functional roles (fig. S8F). The NBD domains of WTN1 were segregated into two distinct clades, with orthologs of NBD I and NBD II dispersed among species within the Pooideae subfamily (fig. S9, S10A-C), suggesting that WTN1 orthologs originated before the LCA of Pooideae. The NBD domains from WTN1-related proteins clustered into two monophyletic groups, and each NBD group was identified throughout the Poaceae, indicating that an ancestral duplication event of the NBD domain probably occurred before the LCA of the Poaceae (fig. S10D). Analysis of the distribution of Rwt4/WTK3 and WTN1 proteins across plant species revealed an intriguing pattern: the absence of Rwt4/WTK3 in a species often coincided with the loss of the WTN1 ortholog (fig. S9, S11). At least five instances of simultaneous losses of both Rwt4/WTK3 and WTN1 were observed (fig. S9), suggesting that the Rwt4/WTK3-WTN1 pair functioned as a cohesive module that was lost entirely during the evolution of Pooideae. Such findings further substantiate the hypothesis that Rwt4/WTK3 and WTN1 co-evolved and acted in concert prior to the LCA of Pooideae.

## Discussion

Tandem kinase proteins (TKPs), arising from the duplication or fusion of kinase domains, represent an emerging class of broad-spectrum disease-resistance proteins widespread in plants (*12-20*). Our interdisciplinary research, including plant genetics, biochemistry, and electrophysiology, has discovered for the first time a class of non-canonical NLRs that are required for the TKP-mediated immune signaling pathway. This unique NLR features dual NB-ARC domains (*23*) and can form calcium-permeable channels upon activation, akin to ZAR1 and Sr35 (*6*, *9*, *11*). However, they differ in the way for activation: Sr35 directly recognizes Avr, whereas WTN1, like ZAR1, relies on auxiliary proteins for indirect Avr detection. Yet, they share a similar but distinct mechanism for Avr detection, “Decoy” vs. “Guard”. ZAR1 teams up with a pseudokinase RKS1 (adaptor) and a PBL2 (sensor) to detect AvrAC modifications on PBL2, while WTN1 cooperates with a TPK harboring two tandem kinase domains with distinct roles - the N-terminal Kin I (including PKF) directly binds Avr as a sensor, and the C-terminal Kin II interacts with NLRs like RKS1 does, transmitting pathogen signals to activate NLR.

The physical closeness between *WTN1* and *Rwt4*/*WTK3* is consistent across the Pooideae subfamily, indicating a conserved genomic arrangement and well-preserved genomic structure that supports their functional interplay. The PKF and Kin I domains have experienced stronger positive selection, aligning with their roles in recognizing PWT4 and potentially the *Bgt* Avr effector, respectively. The concerted losses of the *Rwt4*/*WTK3*-*WTN1* pair in certain species further underscore their co-evolution and joint functional significance predating the LCA of Pooideae, highlighting an ancient strategy for pathogen defense in plants.

Wheat blast and powdery mildew pose major challenges to agriculture, resulting in significant yield reductions worldwide (*24–26*). Identifying their resistance genes and uncovering their mechanisms of action is crucial for breeding and deploying resistant wheat cultivars. Our research, starting with EMS mutagenesis on wheat carrying the *WTK3* gene, discovered *WTN1* as an essential NLR for mediating resistance to powdery mildew, although the specific Avr recognized by WTK3 is yet to be determined. Hence, we turned to study the allelic variant *Rwt4* to decipher the molecular mechanism of TKP-NLR action. Remarkably, we found that both Rwt4 and WTK3 recognize the wheat blast Avr PWT4, and activate WTN1 to form a calcium-permeable channel, thereby triggering immunity. This implies that the *Pm24* (*WTK3*) allele, known for powdery mildew resistance, may also confer resistance to wheat blast, but this hypothesis and the underlying mechanisms deserve further investigation.

Taken together, this study, together with the discovery of Sr62^TK^-Sr62^NLR^ (*27*), uncovers a novel mechanism in plant defense, where TKPs and their cognate NLRs act as “sensor-executor” duos to counteract fungal pathogens in cereal crops. Furthermore, evolutionary analyses reveal a co-evolutionary trajectory of the TKP-NLR module, highlighting their synergistic role in triggering plant immunity.

## Material and methods

### Plant materials and growth conditions

In our previous research, we collected or developed various wheat lines, including the Chinese wheat landraces Hulutou (HLT) and Chiyacao (CYC), a *Pm24* (*WTK3*) gene transgenic line named WTK3-COM4 (created in a Fielder background under the native *WTK3* promoter), and two EMS-induced powdery mildew susceptible *WTK3* mutants (WTK3^W429*^ and WTK3^W478*^) (*17*). For this study, we have generated EMS-induced powdery mildew-susceptible *WTN1* mutants, F_1_ hybrids derived from the crosses of *WTK3* mutants × *WTN1* mutants and *WTN1* mutants × *WTN1* mutants, and a *WTN1* CRISPR/Cas9 knockout line (WTN1-KO) in the WTK3-COM4 T_5_ background. Seeds of the wheat cultivars Jagger and Cadenza were acquired from Professor Caixia Lan at Huazhong Agricultural University, China. Wheat plants were cultivated either at the Gaoyi experimental station in Hebei province or in a controlled greenhouse environment in Beijing for seed propagation. The greenhouse conditions were maintained at a 16 h light/8 h dark cycle with temperatures set to 24/18 °C. For inoculation with *Bgt*, two-week-old seedlings at the two-leaf stage were grown under greenhouse conditions with the same temperature and light regime and a relative humidity of 70%. *N. benthamiana* plants, used for transient expression assays and protein-protein interaction studies, were grown in a greenhouse maintained at 22 °C under long-day conditions (16 h day/8 h night).

### Pathogen inoculation

Wheat seedlings at the two-leaf stage were inoculated with *Bgt* isolate E09 to assess their susceptibility to powdery mildew, following methods previously established (*17*). The evaluation of disease symptoms was conducted at 14 dpi, using an infection type (IT) scale ranging from 0 to 4. This scale is detailed as follows: a score of 0 indicates the absence of visible symptoms; 0; for necrotic flecks, 1 through 4 represent highly resistant, moderately resistant, moderately susceptible, and highly susceptible, respectively. To ensure a consistent supply of the *Bgt* isolate for these experiments, it was propagated and maintained on seedlings of the Fielder cultivar, known for its high susceptibility to powdery mildew. For microscopic observation and documentation of mildew growth, leaves subjected to mildew exposure were stained according to Lu *et al.* (*17*). Observation was made under an Olympus BX-53 microscope (Olympus, Tokyo, Japan). Relevant powdery mildew reaction data are presented in Supplementary Tables S1 and S3.

### Mutagenesis and mutation screening

Mutagenesis and mutation screening of the EMS-induced HLT M_2_ population were performed following the methods in our previous study (*17*). PCR amplification and sequencing of the *WTK3* genes from the powdery mildew-susceptible mutants yielded 16 without a *WTK3* mutation. Further analysis through RNA-Seq, read mapping, and PCR amplification revealed mutations in the *WTN1* gene in five out of the 16 mutants. Details of the primer sequences used are available in Supplementary Table S10.

### VIGS assay

VIGS of *WTN1* was conducted using a recombinant Barley Stripe Mosaic Virus (BSMV) vector, BSMV:*WTN1,* harboring a specific *WTN1* fragment (*WTN1* CDS:1558-1807) (*28*). This vector was used to infect the first set of leaves of 9-day-old HLT seedlings. For controls, HLT seedlings were infected with BSMV:*TaPDS* (the positive control) and BSMV:γ (an empty vector) as the negative control. At 10 dpi, these plants were challenged with the avirulent *Bgt* isolate E09. Powdery mildew reactions were assessed at 14 dpi. Additionally, the third leaves from plants infected with BSMV:γ and BSMV:*WTN1* were harvested and pooled for gene silencing efficiency, with primer sequences detailed in Supplementary Table S10.

### Development of CRISPR/Cas9 genome-edited wheat plants

For the development of CRISPR/Cas9 genome-edited wheat plants, specific target sequences of *WTN1* were synthesized to generate Cas9-sgRNA expression vector constructs, following the protocol reported (*29*). These constructs were introduced into the wheat WTK3-COM4 T_5_ line via *Agrobacterium tumefaciens* strain EHA105, with selection based on PCR and sequencing. The primer sequences for this process are also listed in Supplementary Table S10, and data on powdery mildew reactions are provided in Supplementary Table S3.

### RT-qPCR analysis

Quantitative reverse transcriptase PCR (RT-qPCR) was conducted to evaluate the expression levels of the genes WTK3 and WTN1 in wheat leaves. Total RNA was extracted from wheat leaves using TRIzol reagent (Tiangen, DP419). Reverse transcription was performed using a PrimeScript RT reagent kit with a gDNA Eraser (TaKaRa, RR047). RT-qPCR was performed in a Roche 480 light thermocycler (Roche, Colorado Springs, CO) and analyzed using a SYBR Premix Ex Taq II (TaKaRa, RR820). The primer sequences for *WTN1,* used to evaluate the transcript levels, are detailed in Supplementary Table S10, with the wheat *ACTIN* gene as the internal control for normalization. The comparative threshold 2^−ΔΔCT^ method was used to quantify relative gene expression (*30*). Each sample was analyzed with three replicates. Relevant gene expression data are given in Supplementary Table S2.

### Y2H assay

For the Y2H assay, target coding sequences were cloned into the pGBKT7 and pGADT7 vectors (TaKaRa). These vectors were co-transformed into the Y2HGold yeast strain and cultured on a selective medium devoid of tryptophan (Trp), leucine (Leu), and histidine (His) or lacking Trp, Leu, His, and adenine (Ade), supplemented with X-β-Gal. Images of the colonies were captured at 3 dpi at 30°C. The primer sequences employed for cloning are listed in Supplementary Table S10.

### BiFC and SFLC assays

For BiFC and SFLC assays, WTK3, Rwt4, WTK3-N^1-500^, Rwt4-N^1-502^, WTK3-C^521-893^, Rwt4-C^521-895^, Rwt4 ^Δ PKF^ (Rwt4^146-895^), WTN1 ^Δ 23^ (WTN1^24-1038^) and PWT4 coding sequences were cloned into BiFC vectors pCAMBIA1300-35S-cYFP and pCAMBIA1300-35S-nYFP and SFLC vectors pCAMBIA-cLUC and pCAMBIA-nLUC to generate fusion constructs (*31*). These constructs were transformed into *A. tumefaciens* strain GV3101, which was then cultured overnight in an LB medium. The bacteria were re-suspended in an infiltration buffer (10 mM MgCl_2_, 10 mM methyl ester sulfonate, and 150 μM acetosyringone, at pH 5.6) and incubated for 2 h at 30°C prior to plant infiltration. Constructs were co-transfected into leaf epidermal cells of 5-week-old *N. benthamiana* for protein-protein interaction. Yellow Fluorescent Protein (YFP) fluorescence was observed using a Zeiss LSM710 confocal microscope (Zeiss, Oberkochen, Baden-Württemberg, Germany) 48 h post-infiltration. Additionally, luciferase substrate (Promega, E1605) was sprayed onto the surface of injected *N. benthamiana* leaves, and LUC signals were captured using a cooled CCD imaging apparatus (Berthold, LB985). Both BiFC and SFLC assays were conducted in at least three independent experiments. Primer sequences used for these analyses are provided in Supplementary Table S10.

### *In vivo* pull-down

In the *in vivo* pull-down assay, target coding sequences were cloned into the pGEX-4T-1 vector (GE Healthcare, Chicago, IL) and pET-30a-c vector (Novagen, TB095) for the expression of GST and His-tagged proteins, respectively. The proteins GST-PWT4, GST, His-Rwt4, His-WTK3, His-Rwt4-N^1-502^, His-WTK3-N^1-500^ and His-Rwt4^1-145^ were expressed in *Escherichia coli* Rosetta2 (DE3) (Zomanbio, ZC1211-2). Expression was induced with 0.5 mM isopropyl β-D-1-thiogalactopyranoside (IPTG) at 20°C for 16 h. GST-PWT4 and GST proteins were purified and immobilized on Glutathione Sepharose 4B beads (GE Healthcare, 17-0756). The beads were divided equally into five aliquots and incubated with the same amount of His-Rwt4, His-WTK3, His-Rwt4-N^1-502^, His-WTK3-N^1-500^ and His-Rwt4^1-145^ protein lysate for 2 h at 4°C. The beads were subsequently washed five times with TBS-T buffer [500 mM NaCl, 20 mM Tris-HCl (pH 8.0), 0.1%Tween 20], followed by elution with 100 μL of elution buffer (50 mM Tris-HCl, 10 mM reduced glutathione, pH 8.0). Supernatants were resolved in 10% SDS-PAGE and subjected to immunoblotting using anti-GST (Transgen, HT601-02) and anti-HIS (Transgen, HT501-02) antibodies. The primer sequences used for cloning the target coding sequences into the vectors are provided in Supplementary Table S10.

### Co-IP assay

In the Co-IP assay, target coding sequences were cloned into the pUC57-Ubi-N-3×Flag and pUC57-Ubi-C-3×HA vectors for the expression of fusion proteins with Flag and HA epitope tags in wheat protoplasts, respectively. Fielder protoplasts (3 mL, OD_600_ = 0.8) were transfected with these constructs (Flag-WTK3, 300 μg; WTN1-HA, 100 μg) and incubated overnight in the dark. Total proteins were then extracted from the harvested protoplasts using a specific extraction buffer (150 mM NaCl, 150 mM Tris-HCl, pH 7.5, 1 mM EDTA, 10% glycerol, 10 mM DTT, 0.4% Nonidet-40, 2% PVPP, 1× protease inhibitor cocktail). The coding sequences of WTK3, WTK3-N^1-500^, WTK3-C^521-893^, Rwt4, WTN1^Δ23^ (WTN1^24-1038^) and PWT4 were amplified and then inserted into the pSuper1300-N-3×Flag and pSuper1300-C-3xHA vectors for the expression of fusion proteins with Flag and HA epitope tags in *N. benthamiana*, respectively (*32*). The resulting plasmids were transformed into the *A. tumefaciens* strain GV3101, cultured overnight in LB medium, collected, and resuspended in the infiltration buffer, and incubated for 2 h at 30 °C before infiltration. After 2 d of growth post-infiltration, *N. benthamiana* tissues were harvested, and ground in liquid nitrogen, and the proteins were solubilized with the extraction buffer. The combinations of Flag-WTK3, Flag-WTK3-N, Flag-WTK3-C, Flag-Rwt4, WTN1 ^Δ 23^-HA and PWT4-HA proteins were incubated at 4°C for 2 h to capture Flag-tagged proteins using Flag-Trap magnetic beads (Sigma-Aldrich, M8823). Following incubation, the beads were washed five times with the TBS-T buffer [500 mM NaCl, 20 mM Tris-HCl (pH 8.0), 0.1%Tween 20]. The precipitated proteins were then detected via western blot using anti-HA (MBL, M180-7) or anti-DDDDK (MBL, M185-7) antibodies. The primer sequences used for cloning are listed in Supplementary Table S10.

### BN-PAGE assay

To investigate WTN1 oligomerization, we performed BN-PAGE using the Bis-Tris Native PAGE system (Invitrogen, Carlsbad, CA). The coding sequences of *WTN1*^Δ23^, *PWT4*, and *AvrPm3b2*/*c2* (NCBI MK806483.1) were all synthesized and were cloned into the pUC57-Ubi-C-3×Flag or pUC57-Ubi-C-3×HA vectors under the control of a strong ubiquitin promoter. Transfection involved 100 μg each of WTN1^Δ23^-3×Flag, PWT4-3×HA and AvrPm3b2/c2-3×HA constructs into wheat Jagger (Rwt4/WTN1) protoplasts at an OD_600_ of 0.8, followed by incubation with W5 solution [154 mM NaCl, 125 mM CaCl_2_, 5 mM KCl, 2 mM MES (pH 5.7), 1mM LaCl_3_] for 24 h. Total proteins were extracted with the protein extraction buffer as previously described (*33*), with samples prepared for BN-PAGE by combining protein extracts with NativePAGE^TM^ Sample Buffer (Invitrogen^TM^, BN2003), NativePAGE^TM^, G-250 Sample Additive (Invitrogen^TM^, BN2004), Digitonin (Invitrogen^TM^, BN2006), and distilled water. These samples were then run on Native PAGE 3%-12% Bis-Tris Gels (Invitrogen^TM^, BN1001), with Native Protein Marker (Bioroyee, RZ1828) loaded for protein size estimation. Following electrophoresis, proteins were transferred to a PVDF membrane using NuPAGE Transfer Buffer (Invitrogen^TM^, NP00061) for immunoblot analysis using specific antibodies. The primer sequences used for cloning are detailed in Supplementary Table S10.

### Cell death assay on *N. benthamiana* leaves

The cell death assay in *N. benthamiana* leaves was conducted following established protocols (*34, 35*). Target coding sequences were cloned into the pCAMBIA1300-MAS-nos vector to generate fusion constructs. The resulting plasmids were transformed into *A. tumefaciens* strain GV3101. After culturing overnight in LB medium, the bacteria were resuspended in an infiltration buffer and incubated at 30°C for 2 h prior to leaf infiltration. The leaves were then photographed 72 - 96 h post-infiltration.

### Transient gene expression assay in wheat protoplasts

In the transient gene expression assay conducted in wheat protoplasts, seedlings from various wheat accessions, including Jagger, Cadenza, CYC, HLT, the transgenic T_5_ line WTK3-COM4, the CRISPR/Cas9 knockout T_2_ line WTN1-KO and EMS-induced powdery mildew-susceptible mutants WTK3^W429*^ were grown under controlled greenhouse conditions. Protoplasts were isolated from these plants and transfected as previously described (*36*). The effector genes *PWT4* and *AvrPm3b2/c2* (NCBI MK806483.1), excluding their signal peptides, were synthesized and cloned into the pUC57-Ubi-C-3×HA vector under the control of the ubiquitin promoter. To assess gene expression, co-transfection with the pRTVnMyc vector was performed to facilitate the expression of an LUC reporter gene (*37*). For each experimental setup, 250 µL of wheat protoplasts (OD_600_ = 0.4) were transfected with 10 µg of pRTVnMyc-LUC and either pUC57-Ubi-C-3×HA:PWT4 for the treatment groups or pUC57-Ubi-C-3×HA:AvrPm3b2/c2 for the control groups. Luminescence was measured using the GloMax™ 20/20 (Promega), and relative luminescence values were calculated by normalizing to the control treatment. This process was repeated across three independent experiments to ensure consistency, with results from one representative set being presented. Absolute luciferase values and primer sequences used in this assay are detailed in Supplementary Tables S4 and S10, respectively.

### Electrophysiology

Electrophysiology studies employing the two-electrode voltage clamp (TEVC) technique were conducted to measure ionic currents through *Xenopus* oocytes expressing wild-type or mutant versions of *WTN1*, *WTK3*, *Rwt4*, *PWT4*, and *AvrPm3b2*/*c2*. These cDNAs were cloned into the pGHME2 plasmid, and cRNA was synthesized using T7 polymerase after linearization with *Nhe*I. The procedures involving animals adhered to established standards for laboratory animal care. Oocytes were harvested from adult *X. laevis* frogs under tricaine anesthesia (0.3% 3-aminobenzoic acid ethyl ester), and cRNA injections were optimized to mitigate toxicity associated with the expression of the WTN1 resistosome. A specific mixture of cRNAs for WTN1, WTK3/Rwt4, and PWT4, each at a concentration of 250 ng/μL, was prepared in a volume ratio of 11:11:1 to achieve equal molar expression and injected into the oocytes at a dosage of 36.8 nL per oocyte. Following injection, the oocytes were incubated at 18°C for approximately 4 h in ND-96 buffer (96 mM NaCl, 2.5 mM KCl, 1 mM MgCl_2_, 1.8 mM CaCl_2_, 5 mM HEPES, pH 7.6) supplemented with 10 μg L^-1^ penicillin and 10 μg L^-1^ streptomycin. TEVC measurements were conducted in ND-96 solution, approximately 4 - 5 h post-injection, using water-injected oocytes as controls.

Ionic currents were recorded using the OC-725C oocyte clamp amplifier (Warner Instruments) and a Digidata 1550B digitizer (Axon Instruments) with pClamp software (Molecular Devices). Recording microelectrodes filled with 3 M KCl were used, and the membrane potential was systematically varied to elicit currents, which were then analyzed to assess the functionality of the expressed proteins. The experiments were carried out in a temperature-controlled environment, with data analysis performed using Origin 8.0 software (OriginLab Corporation). The results, including current amplitudes and membrane potentials, were averaged across multiple oocytes and presented as the mean ± SEM. Detailed values and results from these measurements are available in Supplementary Table S6.

### Statistical analysis

Statistical analysis was conducted to evaluate the significance of the results obtained from the transient gene expression assays in wheat protoplasts involving PWT4 and AvrPm3b2/c2, and the TEVC measurements in *Xenopus* oocytes. For the LUC measurements and the transient gene expression assays, one-way analysis of variance (ANOVA) was employed, followed by Tukey′s post hoc test to determine specific differences between groups, with a significant threshold set at *P* < 0.05. Similarly, TEVC measurements from *Xenopus* oocytes were analyzed using one-way ANOVA and Tukeys post -hoc test, but with a more stringent significance level of *P*-value < 0.01. The comprehensive results of these analyses, detailing the statistical significance of observed differences, are presented in Supplementary Tables S5 and S6.

### Evolutionary analyses of Rwt4/WTK3 and WTN1

The evolutionary origins and diversification of Rwt4/WTK3 and WTN1 were systematically explored through a comprehensive analysis involving 122 representative plant species. This cohort included 19 non-flowering plants and 103 flowering plants, with a specific focus on 36 species within the order Poales (Supplementary Table S7). For the identification of Rwt4/WTK3 homologs, a BLASTP search was conducted against the proteomes of these 122 species, using the three kinase domains of *T*. *aestivum* Rwt4/WTK3 (accession No. TraesCS1D01G058900) as queries. The search criterion was set with the *e* cut-off value of 10^-5^ to ensure the retrieval of significantly similar sequences. Subsequent alignments of the significant hits with a set of representative RLKs were performed using MAFFT (*38*, *39*), and large-scale phylogenetic analyses were performed using FastTree (*40*). Sequences closely related to the three kinase domains of Rwt4/WTK3 (PKF, Kin I, and Kin II) were retrieved together with representative RLKs for subsequent analyses. For WTN1 identification, we used the NBD sequences of *T. aestivum* WTN1 (accession No.: TraesCS1D01G059000) as queries to perform PSI-BLAST against 122 proteomes, with an *e*-value of 10^-5^ and 20 iterations (*41*). Significant hits of WTN1 were aligned with NBD sequences from all known *Arabidopsis* NLRs using HMMALIGN in HMMER 3.3.2 (http://hmmer.org/). Phylogenetic analyses were performed using the approximate maximum likelihood method implemented in FastTree (*40*). Significant hits that were closely related to WTN1 NBDs were retrieved as WTN1-related proteins, and WTN1 orthologs were defined according to the species phylogeny. Domain architectures were annotated using HMMSCAN in HMMER 3.3.2 (http://hmmer.org/), with an *e-*value cut-off of 0.01 (Supplementary Tables S8 and S9).

To elucidate the phylogenetic relationships among Rwt4/WTK3 and WTN1 proteins, comprehensive phylogenetic analyses were undertaken. For Rwt4/WTK3 proteins and closely related RLKs, kinase domain sequences were aligned using the localpair algorithm in MAFFT (*39*). Similarly, the NBD sequences of WTN1-related proteins were aligned using the same algorithm for consistency and accuracy. These alignments were further refined using TrimAL for automated trimming and were manually edited (*42*). Phylogenetic trees were generated using the maximum likelihood method as implemented in IQ-TREE 2 (*43*). The best-fit substitution model was assessed using the ModelFinder algorithm (*44*). The robustness of the phylogenetic inferences was evaluated using the UFBoot approach (*45*). The resulting phylogenetic trees were annotated and visualized using the Interactive Tree of Life (iTOL) (*46*).

To dissect the conservation patterns across the three kinase domains and crucial functional sites of Rwt4/WTK3 proteins, logo plots highlighting the catalytic motifs and crucial functional sites within the PKF, Kin I, and Kin II domains were generated using the weblogo tool (*47*). Assessment of the selection pressures acting upon these kinase domains was carried out by estimating the ratio of nonsynonymous to synonymous substation rates (dN/dS). For each kinase clade, coding regions were extracted, and short sequences were excluded, resulting in ten PKF sequences, 26 Kin I sequences, and 31 Kin II sequences. Sequences were aligned based on codons using MUSCLE (*48*). The overall dN/dS ratio was calculated by the MEME algorithm in HyPhy. Sites under positive or negative selection were identified using the FEL algorithm in HyPhy with a *P* value of 0.05. MEME was also employed to identify additional sites under positive selection, providing a comprehensive view of the evolutionary pressures shaping these kinase domains.

## Acknowledgments

We are grateful to Profs. Robert McIntosh, University of Sydney, and Evans Lagudah, CSIRO, Australia, for critical improvement of the manuscript. We thank Prof. Caixia Lan of Huazhong Agricultural University for providing the Jagger and Cadenza seeds and Drs. Brande Wulff, King Abdullah University of Science and Technology, Saudi Arabia, and Peter Dodds, CSIRO, Australia, for valuable discussions.

## Author contributions

P.L., G.Z., J.L. and Z.G. performed experimental procedures and analyzed results. G.W., L.L.D., H.Z., K.Z., Q.W., G.G., M.L., B.H., B.L., W.L., L.D., Y.H., X.C., H.F., and D.Q. contributed to gene expression, protein identification, and greenhouse experiments. Y.C. and C.Y. contributed to field trials and phenotypic analyses. G.Z., M.S., and Y.W. contributed to the Electrophysiology experiment. Z.G. and G.Z.H contributed to the evolutionary analyses. P.L., Y.H.C, G.Z.H. and Z.L. conceived of the idea, designed experiments, analyzed genotypic and phenotypic data, and interpreted results. P.L., H.L., J.Z, G.Z.H., Y.H.C. and Z.L. wrote the manuscript.

## Data and materials availability

All data are available in the main text or the supplementary materials. Alignments and phylogenetic trees generated in this study have been deposited to Mendeley Data: Gong, Zhen; Han, Guan-Zhu (2024), “WTK3/WTN1 pair confer multiple disease resistance in wheat”, and are available at https://data.mendeley.com/preview/9tdwzgh8v7?a=613f1967-6ce8-44e4-b62d-54579e92ec52.

## SUPPLEMENTARY MATERIALS

**Table S1.** Responses to powdery mildew *Bgt* isolate E09 in HLT, *WTK3* mutants, *WTN1* mutants, and the F_1_ hybrids of *WTK3* mutants × *WTN1* mutants, and F_1_ hybrids of *WTN1* mutants × *WTN1* mutants.

**Table S2.** Expression patterns of the *WTN1* in the BSMV:*WTN1* knockdown plants and BSMV-γ infection control.

**Table S3.** Responses to powdery mildew *Bgt* isolate E09 infection in Fielder, WTK3-COM4 T_5_, and WTN1-KO T_1_ plants.

**Table S4.** Raw data for all protoplast measurements presented in the main figures.

**Table S5.** Statistical analysis for cell death comparisons in wheat protoplasts.

**Table S6.** Raw data of the electrophysiology data shown in Fig. 4 and Fig. S7.

**Table S7.** Information about the plant species used in this study.

**Table S8.** Domain architectures of WTK3 proteins identified in plant proteomes.

**Table S9.** Domain architectures of WTN1 proteins identified in plant proteomes.

**Table S10.** List of primers used in this study.

**Fig. S1.**
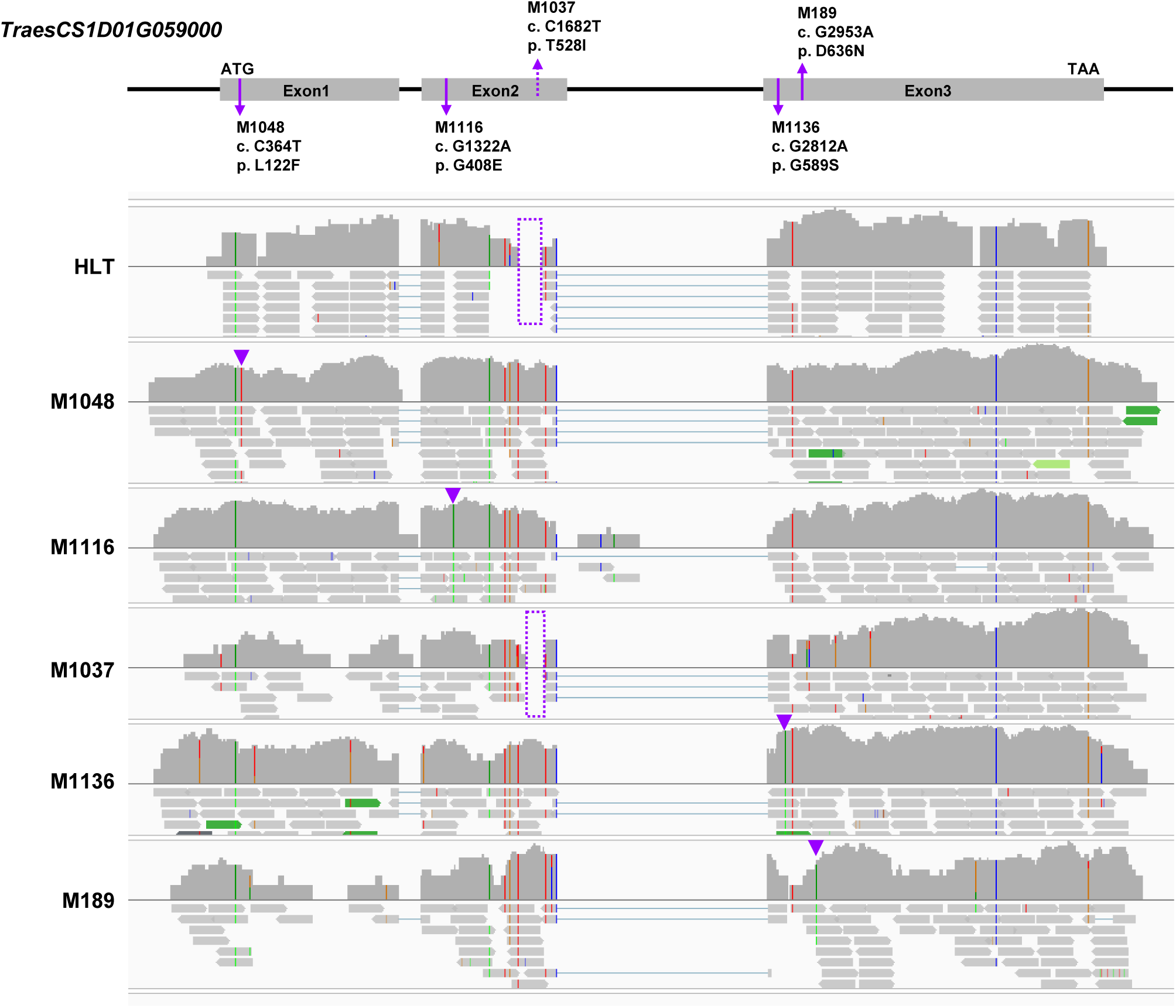
Mapping of RNA-Seq reads from HLT and *WTN1* mutants. RNA-Seq reads from HLT and five EMS-induced HLT mutants susceptible to powdery mildew revealed mutations in an *NLR* gene (*TraesCS1D01G059000*), located 114 kb adjacent to *WTK3* (*TraesCS1D01G058900*) on chromosome 1DS. No sequence variation was observed on *WTK3* in the five mutants. The RNA-Seq reads were aligned to the chromosome 1DS of the Chinese Spring genome RefSeq V1.0. Bases matching the reference genome are shown in gray, while mismatches are colored according to their nucleotide type: A, T, C, and G in green, red, blue, and orange, respectively. Mutation sites in the mutants are indicated by purple triangles, and areas with no reads alignment to the reference genome are highlighted with purple triangles. The mutation M1037 was validated by PCR.

**Fig. S2.**
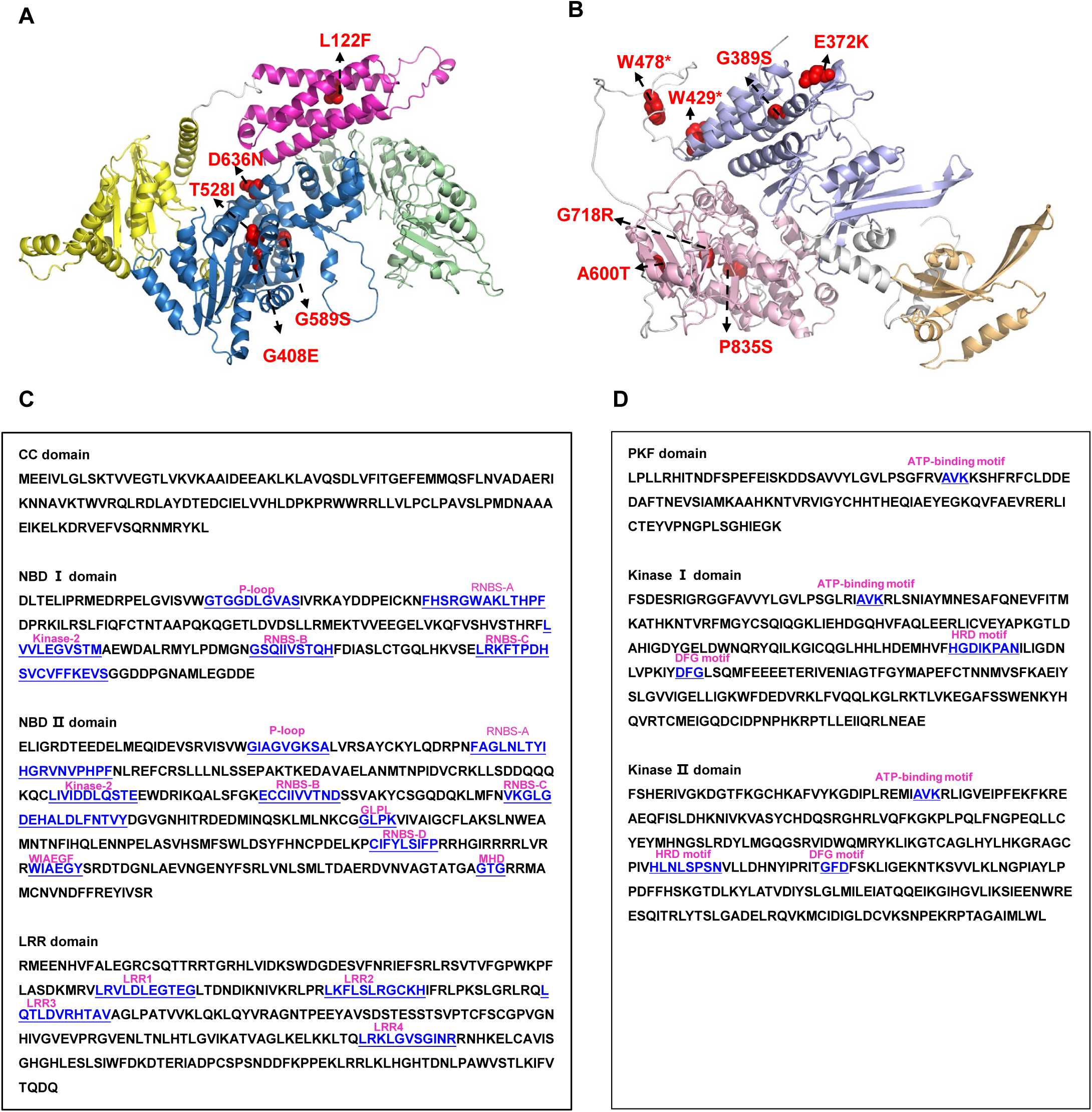
Predicted protein structures and amino acid sequences of WTN1 and WTK3. **(A)** Prediction of WTN1 protein structure by AlphaFold2 showcases the coiled-coil (CC), nucleotide-binding domain I (NBD I), NBD II, and leucine-rich repeat (LRR) domains in magentas, light orange, sky blue and pale green, respectively. The five non-synonymous mutations identified in WTN1 mutants are indicated in red on spherical representations of the residues. **(B)** Predicted structure of WTK3 using AlphaFold2. The PKF, Kin I and Kin II domains are highlighted in light orange, light blue, and light pink, respectively. Two nonsense and five nonsynonymous mutations in WTK3 mutants are indicated in red on spherical representations of the residues. **(C)** Conserved motifs characteristic of the WTN1_NBD I, WTN1_NBD II and WTN1_LRR domains are highlighted in blue characters, and the corresponding names of motifs are indicated with red letters. **(D)** Conserved motifs characteristic of the WTK3_PKF, WTK3_Kin I and WTK3_Kin II domains are highlighted in blue characters, and the corresponding names of motifs are indicated with red letters.

**Fig. S3.**
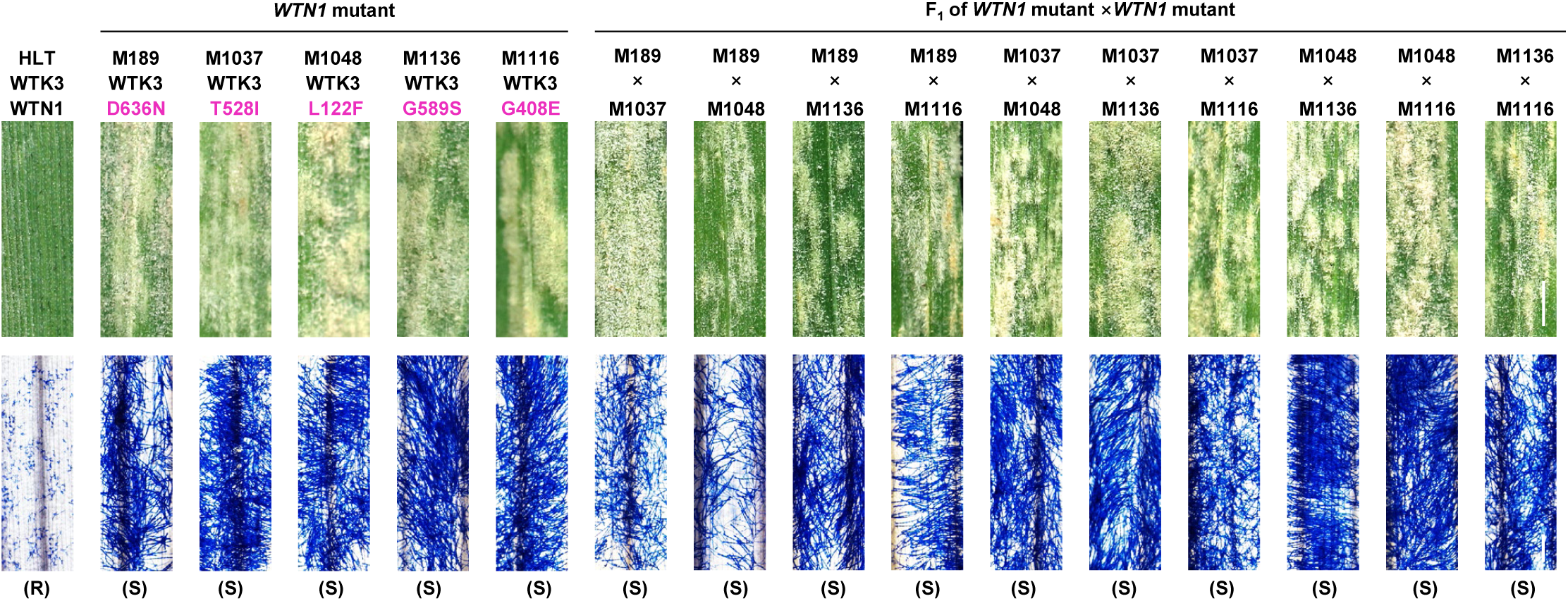
Phenotypes of the F_1_ hybrids from half diallel crosses of *WTN1* mutants. Infection phenotypes of the wild-type HLT, susceptible mutants of *WTN1,* and their F_1_ hybrids from half diallel crosses, inoculated with *Blumeria graminis* f. sp. *tritici* (*Bgt*) isolate E09. Photographs of representative leaves at 14 d post-inoculation (dpi). Scale bar, 0.3 cm. Trypan blue staining of the leaves infected with *Bgt* isolate E09 at 14 dpi to visualize fungal structures. Scale bar, 200 μm. The wild type genotypes of *WTK3* and *WTN1* in the HLT mutants are marked in black, the mutant genotypes of *WTN1* in the HLT mutants are marked in red.

**Fig. S4.**
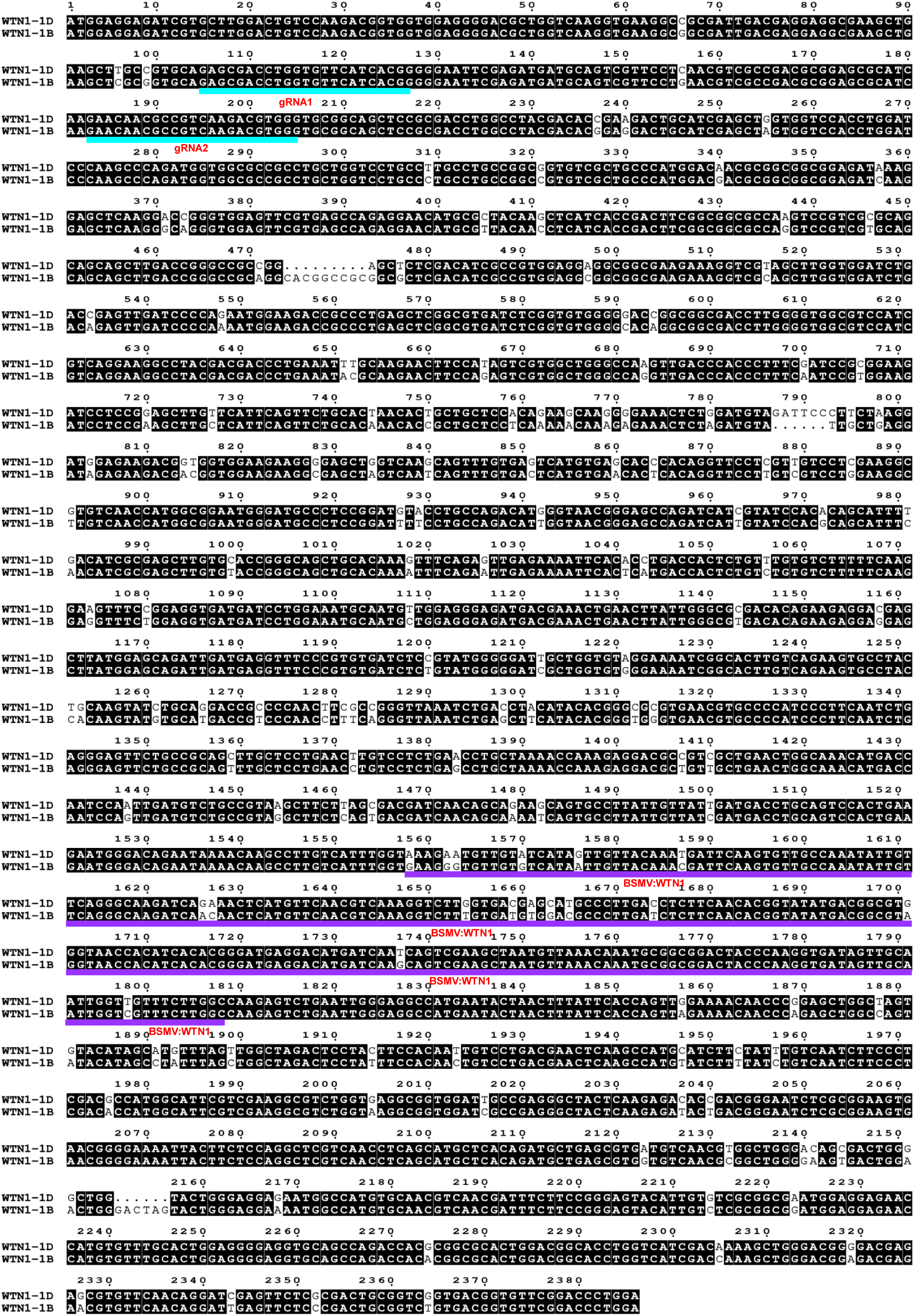
Comparison of the coding sequence (CDS) of *WTN1* and *WTN1-1B* homoeolog. Two single gRNAs (gRNA1 and gRNA2) targeting the 5’-terminal of *WTN1* (*WTN1-1D*) and *WTN1-1B* are highlighted with blue lines. Sequences targeted for virus-induced gene silencing (VIGS) of *WTN1* are highlighted in purple.

**Fig. S5.**
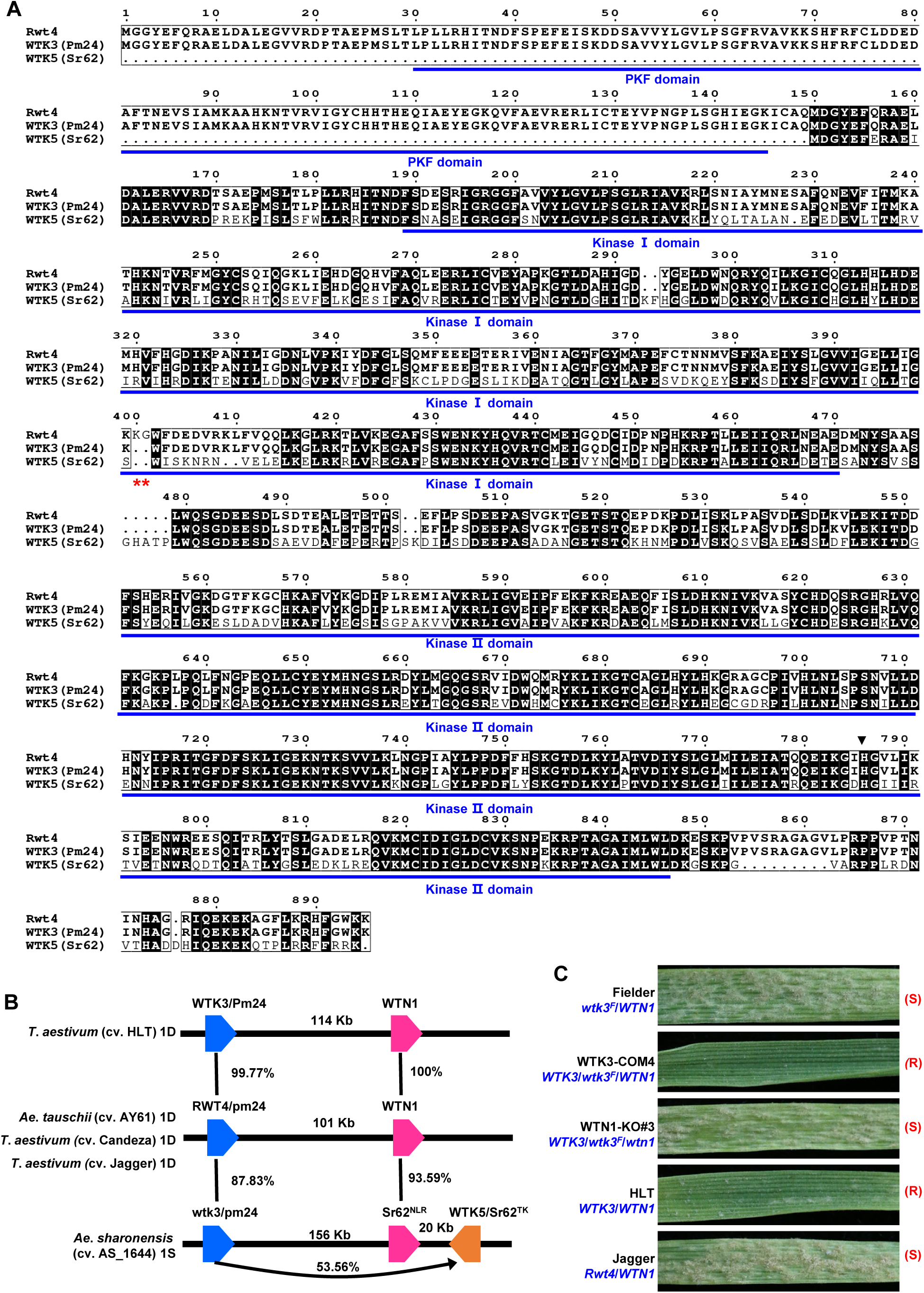
Comparative analysis of WTK proteins. **(A)** Comparison of amino acid sequences of Rwt4, WTK3 (Pm24) and WTK5 (Sr62) using MEGA7. The PKF, Kinase I and Kinase II domains highlighted with blue lines. The amino acid sequence difference between Rwt4 and WTK3 was marked with red asterisks. **(B)** Genomic organization and amino acid sequence comparisons of *WTK3* (*Pm24*), *Rwt4*, *WTK5* (*Sr62*) and *WTN1* in different accessions, including common wheat HLT, Candeza, Jagger and *Aegilops tauschii* AY61, and *Ae*. *Sharonensis* AS_1644. **(C)** Infection phenotypes of Fielder, WTK3-COM4, WTN1-KO#3, HLT, and Jagger inoculated with *B*. *graminis* f. sp. *tritici* isolate E09. Representative leaves were photographed at 14 days post-inoculation (dpi). Genotypes are described in blue. R indicated resistance to powdery mildew, S indicated susceptibility to powdery mildew.

**Fig. S6.**
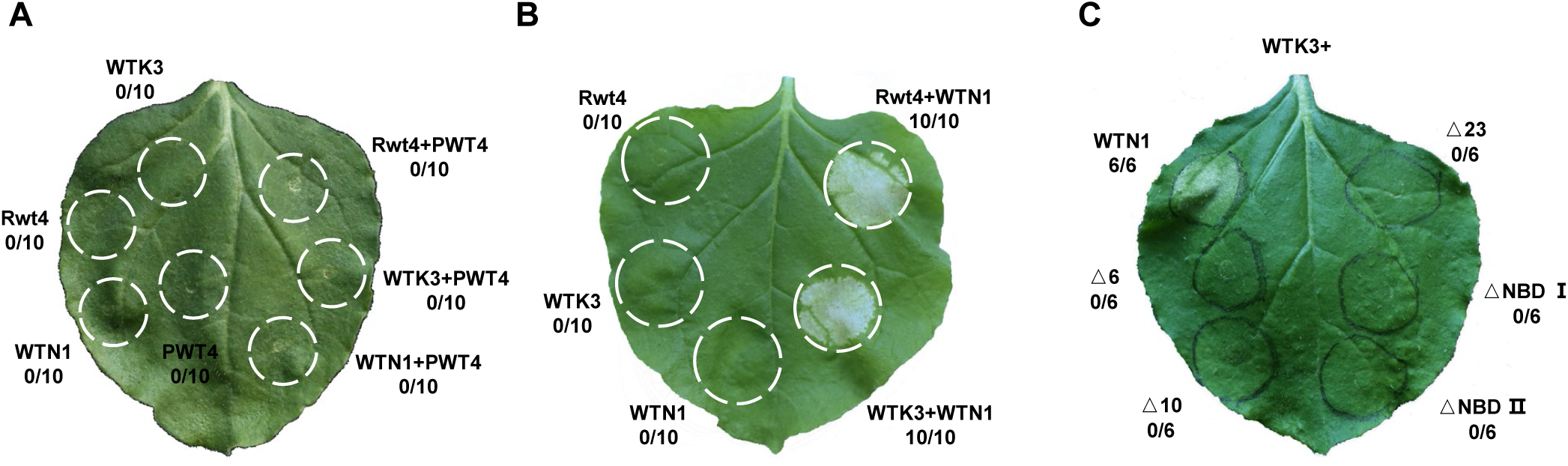
Cell death assays of PWT4, Rwt4/WTK3, WTN1 in *Nicotiana benthamiana* leaves. **(A)** Co-expression of PWT4 with Rwt4, WTK3 and WTN1 in *N*. *benthamiana* leaves. **(B)** Co-expression of WTN1 with Rwt4 and WTK3 in *N*. *benthamiana* leaves. **(C)** Cell death response in *N*. *benthamiana* triggered by co-expression of WTK3 with full length WTN1, N-terminally truncated WTN1 (WTN1^Δ6^, WTN1^Δ10^, and WTN1^Δ23^), WTN1 lacking NBD I (WTN1^ΔNBD I^) and WTN1 without NBD II (WTN1^ΔNBD II^).

**Fig. S7.**
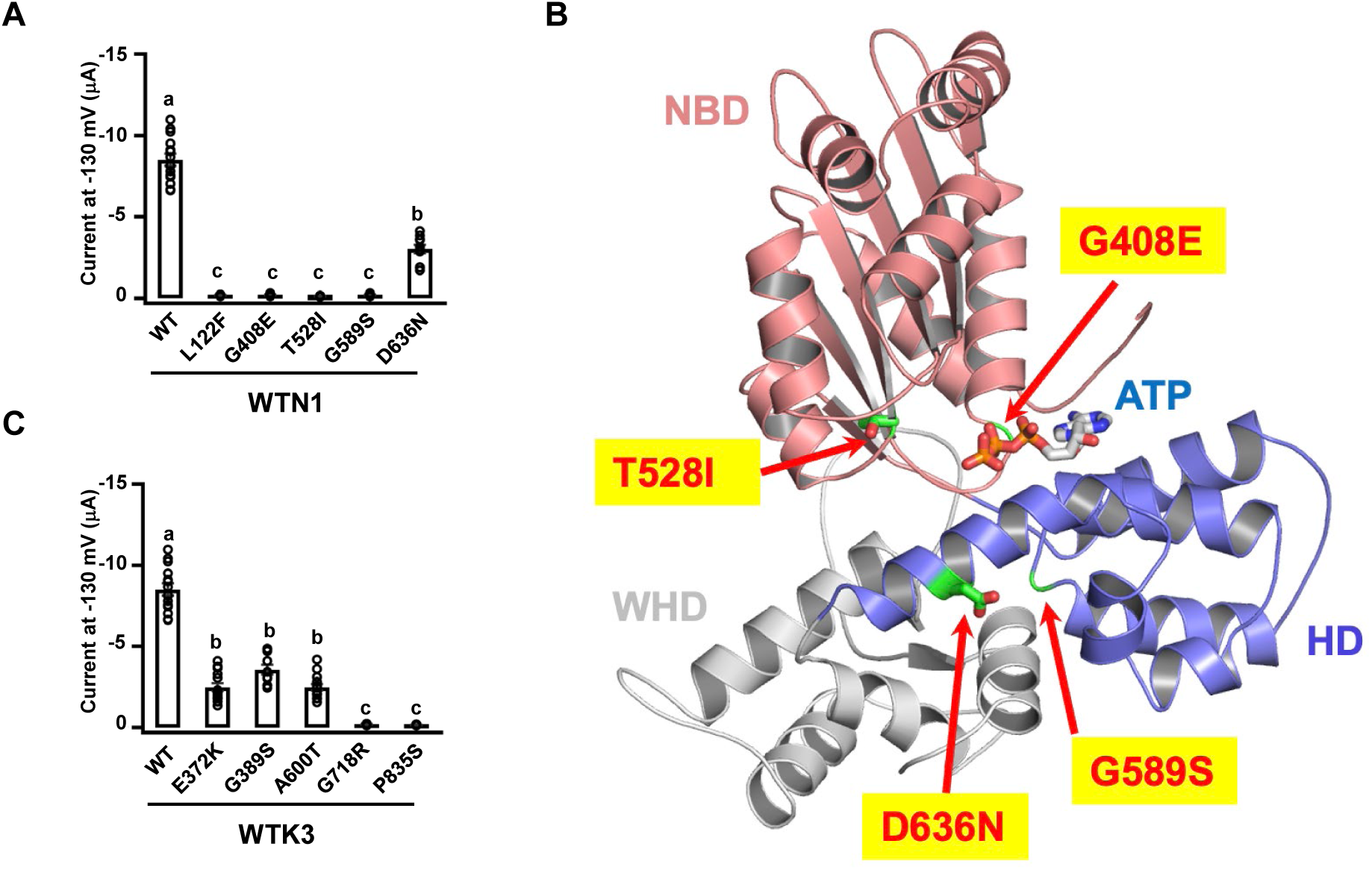
Electrophysiology assays of *WTN1* and *WTK3* mutants and structure prediction of the mutations in WTN1_NBD II. (A) Mutational analysis of WTN1, including L122F, G408E, T528I, G589S and D636N. (B) Structural prediction of WTN1_NBD II, based on AlphaFold2, is depicted in a ribbon diagram. The domains are color-coded: NBD in salmon, HD in light blue, and WHD in grey. An ATP molecule, modeled after the ZAR1 structure (PDB ID: 6j5t), is shown as sticks. Four mutations generated through EMS-generated (Ethyl methane sulfonate) mutagenesis -G408E, T528I, G589S, and D636N-are highlighted within this structure, providing a molecular basis for understanding the functional impacts of these mutations on protein activity. (C) Mutational analysis of WTK3, including E372K, G389S, A600T, G718R and P835S. The current amplitudes measured at -130 mV are shown. Data are presented as means ± SEM, n ≥ 8. Different letters denote the significance tested by one-way analysis of variance (ANOVA) and Tukeys post -hoc test at *P* < 0.01. Exact *P* values are provided in Supplementary Table 6.

**Fig. S8.**
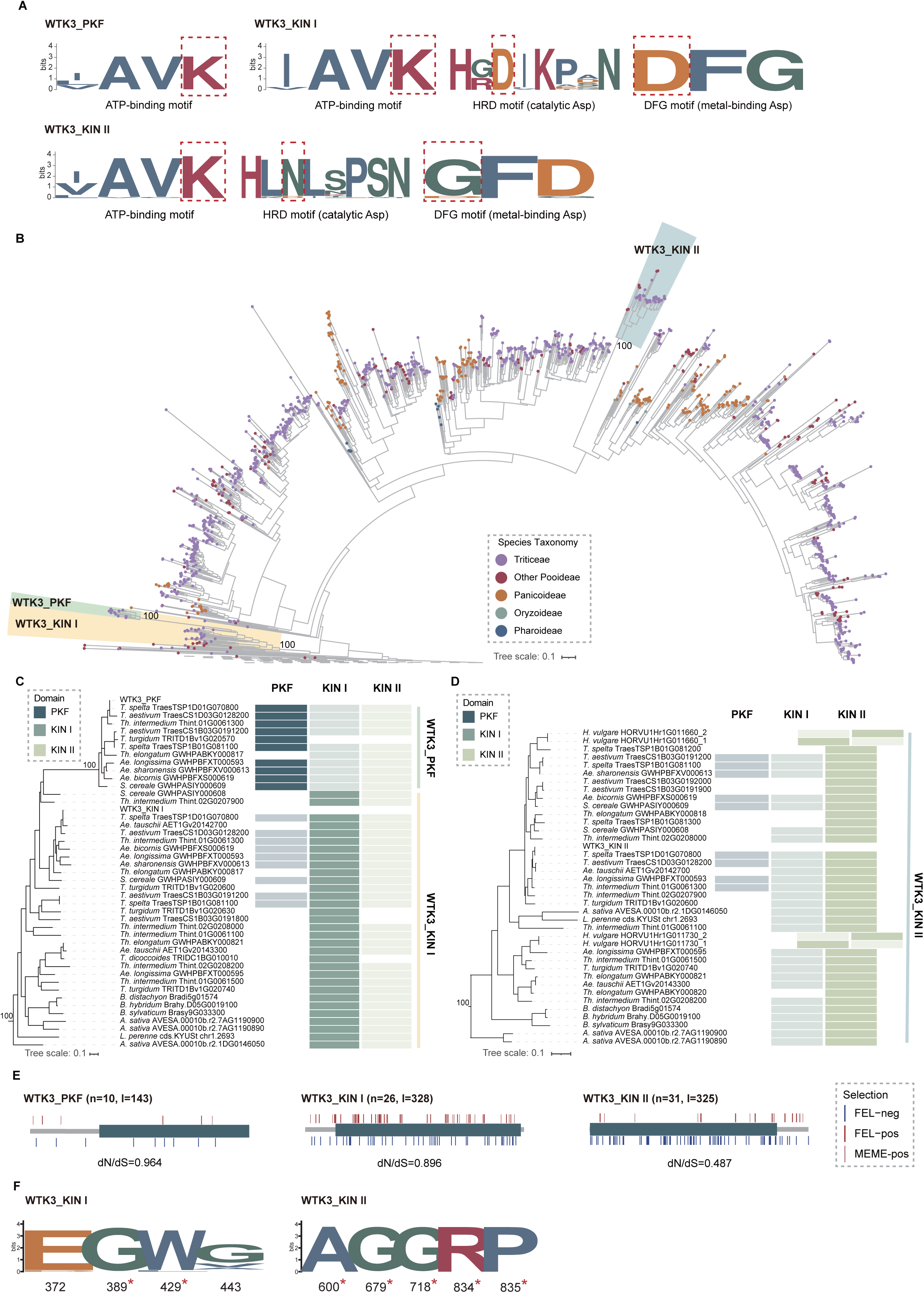
Phylogenetic analysis of evolutionary dynamics of WTK3 kinases. **(A)** Conservation patterns of catalytic motifs across the three kinase domains are visualized through logo plots, emphasizing key catalytic sites with red dashed rectangles. (**B**) The phylogenetic tree, constructed from kinase domain sequences, delineates the relationship among WTK3 kinases and representative RLKs, with WTK3_PKF, WTK3_Kin I, and WTK3_Kin II clades distinguished by green, yellow, and blue, respectively. Ultrafast Bootstrap (UFBoot) support values are provided at clade nodes, and species are represented by solid circles colored according to their taxonomic classification. (**C** and **D**) Detailed phylogenetic trees for WTK3_PKF/Kin I **(C)** and WTK3_Kin II **(D)** sequences featuring the domain architectures adjacent to each protein. Various domains are depicted in distinct rectangles with different colors, with kinase domain analyzed highlighted in a darker shade. *Hordeum vulgare* HORVU1Hr1G011660_1/2 and *H. vulgare* HORVU1Hr1G011730_1/2 indicate these proteins harbor two kinase copies within KIN II clade. **(E)** Selection pressure acting on the WTK3 kinases is explored, presenting the ratio of nonsynonymous (dN) to synonymous (dS) substitutions below each domain. The number of sequences (n) and the alignment length (l) are noted, with short vertical lines marking sites under positive selection (red for FEL, pink for MEME) and negative selection (steel blue for FEL). This selection pressure insight reveals regions of the kinase domains that have undergone evolutionary adaptation, potentially reflecting functional diversification or constraints within the WTK kinase clades. **(F)** Conservation patterns of crucial functional sites analyzed in this study are visualized through logo plots, emphasizing sites under negative selection with red asterisks.

**Fig. S9.**
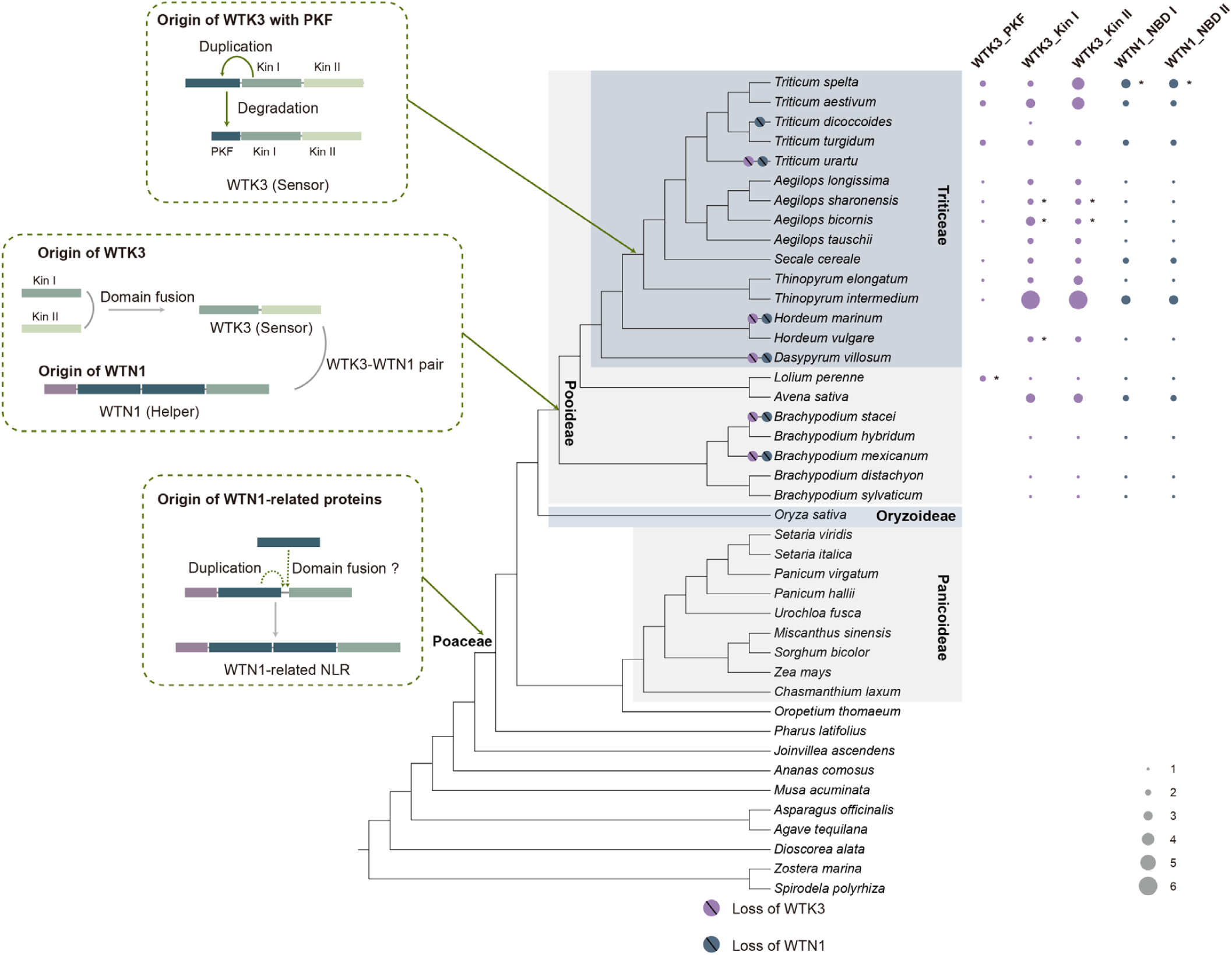
Evolutionary origin and diversification of the WTK3/WTN1 pair in monocots. The phylogenetic distribution and evolutionary development of WTK3 and WTN1 proteins across 42 monocot species are illustrated. WTK3 proteins are categorized by their three kinase domains: WTK3_PKF, WTK3_Kin I, and WTK3_Kin II. WTN1 proteins are differentiated by two NBD domains: WTN1_NBD I and WTN1_NBD II. The prevalence of each domain in the species studied is depicted using solid circles - purple for WTK3 and steel blue for WTN1 – with the circle size indicating the domain copy number. As asterisk signifies proteins identified through tBLASTn, suggesting their presence beyond genome annotated proteomes. The phylogeny of 42 monocots is inferred from literature (*49*) and TimeTree (https://timetree.org/), with circles marked by slashes indicating the evolutionary losses of either WTK3 or WTN1. The boxes on the right side of the phylogeny represent the possible evolutionary scenarios of the WTK3-WTN1 pair. First, WTN1-related NLRs, characterized by dual NBD domains, emerged from NBD duplication and fusion events prior to the last common ancestor (LCA) of the Poaceae family. Subsequently, WTK3 arose through distinct kinase domain fusion before the LCA of Pooideae subfamily, incorporating WTN1 as a pair. This marked the beginning of a co-evolutionary relationship between WTK3 and WTN1 throughout Pooideae evolution, highlighting their intertwined evolutionary paths and functional interdependence.

**Fig. S10.**
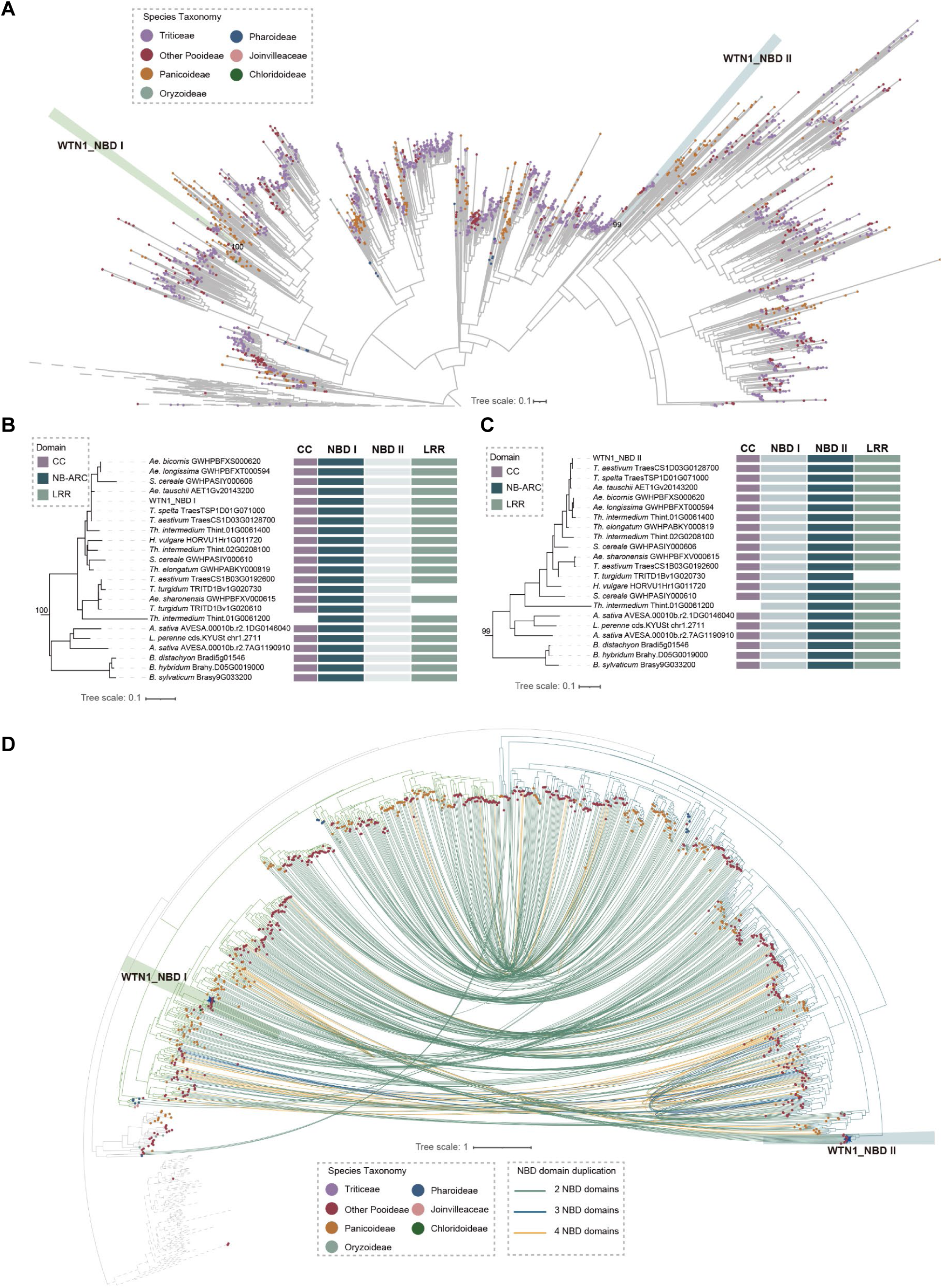
Evolutionary history of WTN1 proteins. **(A)** Phylogenetic relationship among the WTN1 proteins and other NLRs. The phylogeny was reconstructed based on the NBD domain. Two WTN1 NBD clades are accentuated in green and blue. Ultrafast Bootstrap (UFBoot) support values are provided at each clade node, with species indicated by colored circles at the tips, classified by host taxonomy. (**B** and **C**) Detailed phylogenetic analyses for WTN1_NBD I **(B)** and WTN1_NBD II sequences **(C)**, accompanied by illustrations of domain architectures adjacent to each protein. The NBD domain analyzed are highlighted in a darker color. **(D)** Phylogenetic relationship of NBDs of WTN1-related proteins. The phylogeny was reconstructed based on the NBD domains of WTN1-related proteins across representative species, with WTN1_NBD I and WTN1_NBD II clades highlighted in green and blue, respectively. *Triticum aestivum* WTN1 NBDs are marked with blue pentagrams, and NBDs from the same protein are interconnected by colored lines, indicating the presence of multiple NBD domains.

**Fig. S11.**
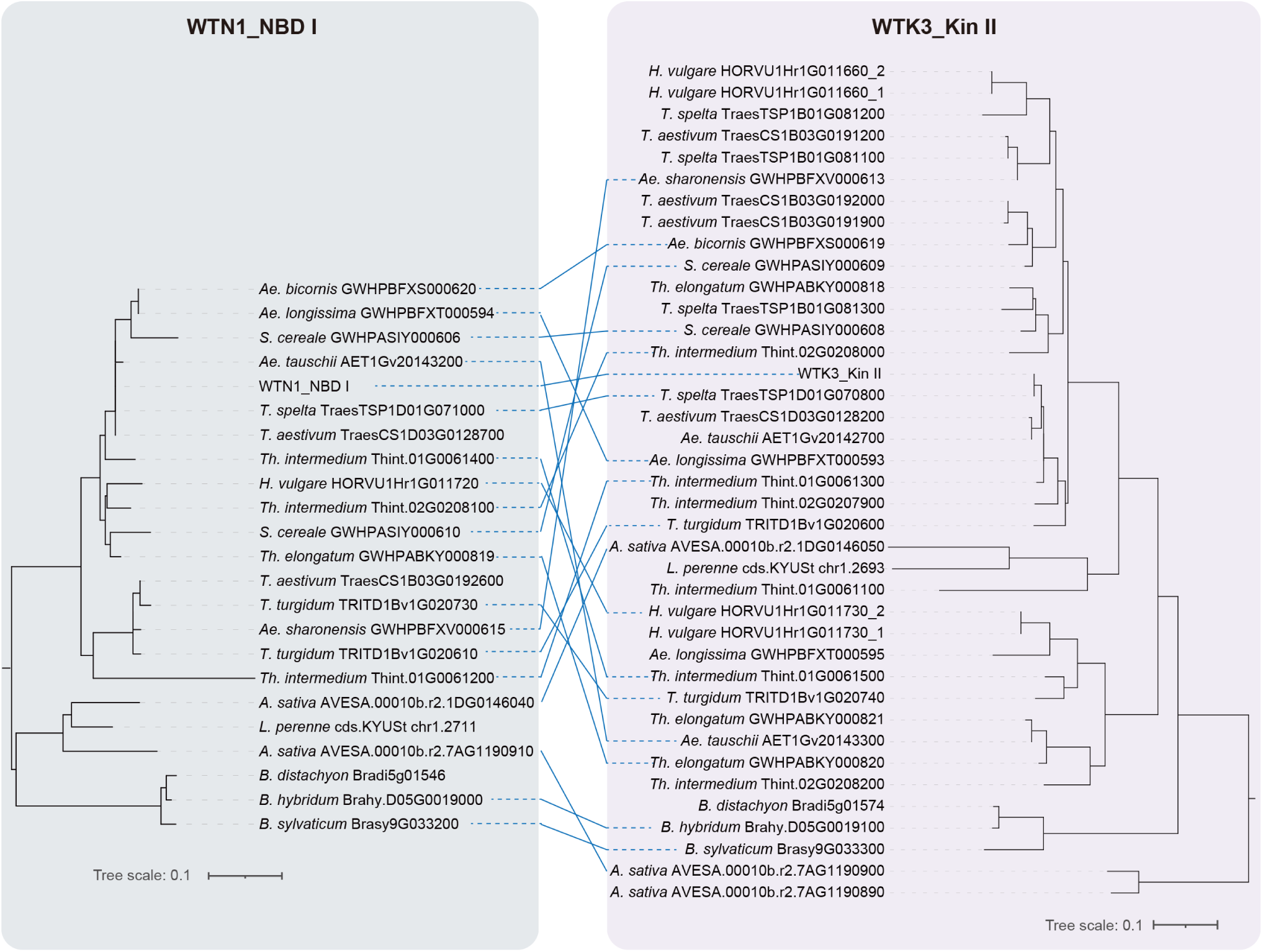
Pairwise association between WTN1 and WTK3 proteins. The phylogenies of WTN1_NBD I and WTK3_Kin II are used as representative frameworks to illustrate the pairwise connections between WTN1 and adjacent WTK3 proteins, highlighting the intricate relationships and potential functional associations between these protein families through blue connecting lines.

